# The *Venturia inaequalis* effector repertoire is expressed in waves and is dominated by expanded families with predicted structural similarity to avirulence proteins from other plant-pathogenic fungi

**DOI:** 10.1101/2022.03.22.482717

**Authors:** Mercedes Rocafort, Joanna K. Bowen, Berit Hassing, Murray P. Cox, Brogan McGreal, Silvia de la Rosa, Kim M. Plummer, Rosie E. Bradshaw, Carl H. Mesarich

**Author notes:** Corresponding author: Carl H. Mesarich. Postal address: School of Agriculture and Environment, Massey University, Private Bag 11222, Palmerston North 4442, New Zealand.

## Abstract

**Background:** Scab, caused by the biotrophic fungus *Venturia inaequalis*, is the most economically important disease of apples worldwide. During infection, *V. inaequalis* occupies the subcuticular environment, where it secretes virulence factors, termed effectors, to promote host colonization. Consistent with other plant-pathogenic fungi, many of these effectors are expected to be non-enzymatic proteins, some of which can be recognized by corresponding host resistance proteins to activate plant defences, thus acting as avirulence determinants. To develop durable control strategies against scab, a better understanding of the roles that these effector proteins play in promoting subcuticular growth by *V. inaequalis*, as well as in activating, suppressing or circumventing resistance protein-mediated defences in apple, is required.

**Results:** We generated the first comprehensive RNA-seq transcriptome of *V. inaequalis* during colonization of apple. Analysis of this transcriptome revealed five temporal waves of gene expression that peaked during early, mid or mid-late infection. While the number of genes encoding secreted, non-enzymatic proteinaceous effector candidates (ECs) varied in each wave, most belonged to waves that peaked in expression during mid-late infection. Spectral clustering based on sequence similarity determined that the majority of ECs belonged to expanded protein families. To gain insights into function, the tertiary structures of ECs were predicted using AlphaFold2. Strikingly, despite an absence of sequence similarity, many ECs were predicted to have structural similarity to avirulence proteins from other plant-pathogenic fungi, including members of the MAX, LARS, ToxA and FOLD effector families. In addition, several other ECs, including an EC family with sequence similarity to the AvrLm6 avirulence effector from *Leptosphaeria maculans*, were predicted to adopt a KP6-like fold. Thus, proteins with a KP6-like fold represent another structural family of effectors shared among plant-pathogenic fungi.

**Conclusions:** Our study reveals the transcriptomic profile underpinning subcuticular growth by *V. inaequalis* and provides an enriched list of ECs that can be investigated for roles in virulence and avirulence. Furthermore, our study supports the idea that numerous sequence-unrelated effectors across plant-pathogenic fungi share common structural folds. In doing so, our study gives weight to the hypothesis that many fungal effectors evolved from ancestral genes through duplication, followed by sequence diversification, to produce sequence-unrelated but structurally similar proteins.

## Background

Fungal pathogens are responsible for some of the most devastating diseases of crop plants worldwide, causing large economic losses, and threatening both food production and security [1]. Resistance to these pathogens is largely governed by the plant immune system, and is based on the recognition of invasion patterns (IPs) by plant immune receptors [2]. At the plant cell surface, many of these immune receptors are pattern recognition receptors (PRRs) of the receptor-like protein (RLP) or receptor-like kinase (RLK) classes, which recognize conserved IPs known as microbe-associated molecular patterns (MAMPs) [3]. Following the recognition of these MAMPs, plant defence responses that slow or halt growth of the fungal pathogen are initiated [3]. To circumvent or suppress these defence responses, plant-pathogenic fungi must secrete a collection of virulence factors, termed effectors, to the plant– pathogen interface, a subset of which may be taken up intracellularly. These effectors are predominantly non-enzymatic proteins, but can also be enzymes, secondary metabolites and small RNAs [4–7]. In some cases, however, effectors can be recognized as IPs by extracellular PRRs or intracellular nucleotide-binding leucine-rich repeat (NLR) immune receptors, collectively referred to as qualitative resistance (R) proteins, to activate plant defences [3, 8]. Such effectors are termed avirulence (Avr) effectors because their recognition typically renders the fungal pathogen unable to cause disease.

Scab (or black spot), caused by *Venturia inaequalis*, is the most economically important disease of apple (*Malus* x *domestica*) worldwide [9, 10]. Under favourable conditions, this disease can render the fruit unmarketable and cause a yield reduction of up to 70% [10]. During biotrophic host colonization, *V. inaequalis* exclusively colonizes the subcuticular environment without penetrating the underlying plant epidermal cells [9, 11, 12]. It is here that the fungus develops specialized infection structures, known as stromata and runner hyphae [9, 11]. Stromata give rise to asexual conidia, but are also likely required for nutrient acquisition and effector secretion [9, 11, 13]. Runner hyphae, on the other hand, enable the fungus to radiate out from the initial site of host penetration, acting as a base from which additional stromata can be differentiated [11]. At the end of the season, in autumn, *V. inaequalis* switches to saprobic growth inside fallen leaves, where it undergoes sexual reproduction [9].

Scab disease is largely controlled through fungicides, which greatly accelerate the development of fungicide resistance in *V. inaequalis* [14, 15]. Scab-resistant apple cultivars, developed through the incorporation of *R* genes, represent a more sustainable disease control option [16]. However, races of *V. inaequalis* that can overcome resistance in apple mediated by single *R* genes have been identified [16, 17]. Therefore, it is likely that multiple ‘unbroken’ *R* genes, perhaps together with quantitative trait loci (QTL), will need to be stacked in each of these cultivars to provide durable disease resistance [16, 17]. For this to be successful, prior knowledge of the molecular mechanisms used by the fungus to overcome *R* gene-mediated resistance, including an understanding of how these mechanisms impact effector function and pathogen fitness, will be required. So far, however, there have been no publications describing the cloning of *Avr* effector genes from *V. inaequalis.* Crucially, the genomes of multiple *V. inaequalis* isolates have been sequenced [18–23] and bioinformatic studies have identified a large catalogue of secreted, non-enzymatic proteinaceous effector candidates (ECs) from which candidate Avr effectors can be identified [19]. Most of these ECs belong to expanded (sequence-related) families [19]; however, the driving force behind the expansion of these families is not yet known. Nevertheless, many *EC* genes are located next to repetitive sequences in the genome of *V. inaequalis* and, therefore, it is possible that transposition is one of the mechanisms driving this expansion [19]. Examples where *EC* genes of *V. inaequalis* are located next to repetitive sequences include members of the *AvrLm6-like* family, which encode proteins with sequence similarity to AvrLm6 [13], an Avr effector from the fungal pathogen *Leptosphaeria maculans* (blackleg of canola) [24], as well as members of the *Ave1-like* family [19], which encode proteins with sequence similarity to Ave1, an antimicrobial Avr effector from the fungal pathogen *Verticillium dahliae* (Verticillium wilt disease) [25, 26].

To develop durable control strategies against scab disease, a better understanding of the roles that effectors play in promoting subcuticular growth by *V. inaequalis* is also required. To date, subcuticular growth has been largely understudied, even though it is exhibited by many plant-pathogenic fungi, including other crop-infecting members of the *Venturia* genus [11, 27–29], as well as, for example, *Rhynchosporium* (scald disease of graminaceous plants) [30, 31] and *Diplocarpon* (e.g. rose black spot) [32, 33]. In recent years, host colonization by plant-pathogenic fungi has been studied by transcriptomic analysis [34–36]. However, comprehensive transcriptomic studies focusing on the subcuticular parasitic strategy are not yet available. Indeed, while previous expression data from interactions between *V. inaequalis* and susceptible apple have been published [19, 37], these data are only based on a limited number of infection time points with no biological replicates.

In this study, we provide the first comprehensive transcriptomic analysis of *V. inaequalis* during colonization of susceptible apple and identify infection-related temporal expression waves associated with *EC* genes of this fungus. Using recent advances in *de novo* protein folding algorithms, we also show that the EC repertoire of *V. inaequalis* is dominated by expanded families with predicted structural similarity to Avr proteins from other plant-pathogenic fungi. Collectively, this study furthers our understanding of subcuticular growth by *V. inaequalis* and provides an enriched list of ECs that can be investigated for potential roles in virulence and avirulence.

## Results

### The different stages of host infection by *V. inaequalis* observed by bright-field microscopy display distinct gene expression profiles

To investigate changes in *V. inaequalis* gene expression during host colonization and relative to growth in culture, we set up an infection time course involving detached leaves of susceptible apple cultivar *M.* x *domestica* ‘Royal Gala’ and compared it to growth of the fungus on cellophane membranes overlying potato dextrose agar (PDA). Here, six *in planta* time points (12 and 24 hours post-inoculation [hpi], as well as 2, 3, 5 and 7 days post-inoculation [dpi]) and one in culture time point (7 dpi) were used.

Analysis of leaf material from the infection time course by bright-field microscopy revealed that, at 12 hpi, conidia of *V. inaequalis* had germinated and formed appressoria on the leaf surface (**Fig. 1A**). At 24 hpi, primary hyphae had developed, indicating that colonization of the subcuticular environment was underway. Then, by 2 and 3 dpi, stromata had differentiated from primary hyphae and, in many cases, these stromata had undergone a rapid expansion in size through non-polar division (**Fig. 1A**). Subcuticular runner hyphae had also started to radiate out from stromata (**Fig. 1A**). By 5 dpi, fungal biomass had accumulated extensively in the subcuticular environment and additional stromata had started to develop from runner hyphae (**Fig. 1A**). Often, these stromata had formed conidiophores, from which conidia had developed (**Fig. 1A**). Finally, at 7 dpi, conidia of *V. inaequalis* had started to rupture through the plant cuticle (**Fig. 1A**) and the first macroscopic olive-brown lesions, indicative of heavy sporulation, were apparent. This represented the end of the biotrophic infection stage, as detached apple leaves had started to decay after this time point.

**Fig. 1.**
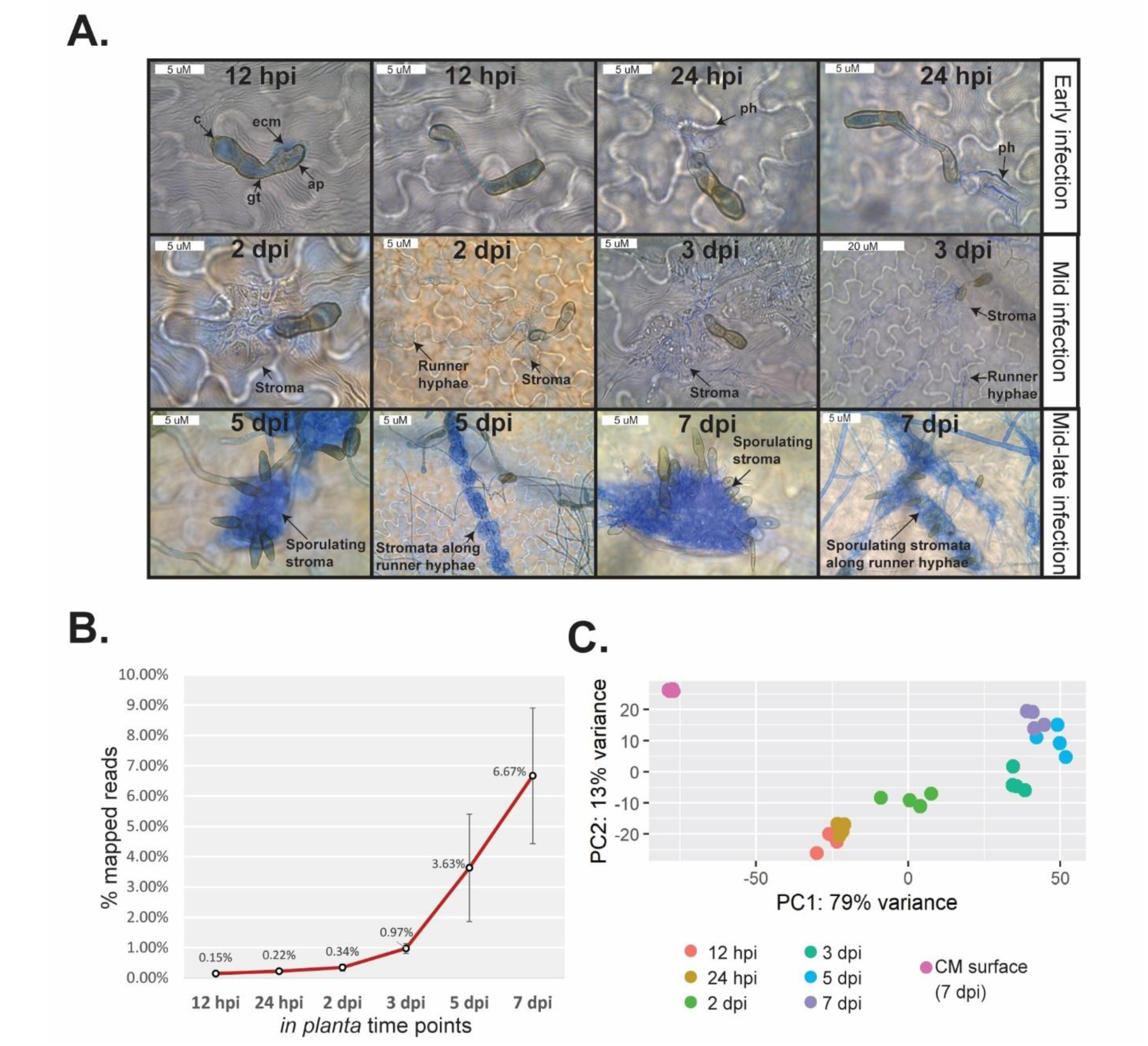
Microscopic and transcriptomic profile of *Venturia inaequalis* during infection of detached leaves from susceptible apple cultivar ‘Royal Gala’. **A.** Microscopic evaluation of *V. inaequalis* during colonization of apple leaves. Infection structures observed by bright-field microscopy were stained with aniline blue. Infected leaves are representative of material used in the RNA-seq transcriptome sequencing experiment. c, conidium; gt, germ tube; ap, appressorium; ecm: extracellular matrix; ph: primary hyphae. **B.** Percentage (%) of paired RNA-seq reads mapped to the *V. inaequalis* genome relative to the total RNA-seq reads sequenced. Error bars represent standard deviation across four biological replicates. **C.** Principal component analysis (PCA) of RNA-seq data from *V. inaequalis* during colonization of apple leaves and in culture on the surface of cellophane membranes (CMs) overlaying potato dextrose agar. Four biological replicates per time point are shown. hpi: hours post-inoculation; dpi: days post-inoculation.

Inspection of the RNA-seq data revealed that the percentage of reads mapping to the *V. inaequalis* genome [19] increased as the infection time course progressed (**Fig. 1B**, **Additional file 1: Table S1**). Furthermore, the biological replicates clustered robustly within time points and a clear distinction between the early and mid-late infection stages, as well as between the *in planta* and in culture growth conditions, was observed (**Fig. 1C**).

### Genes of *V. inaequalis* are expressed in temporal waves during infection of apple leaves

We set out to identify which genes of *V. inaequalis* are up-regulated during infection of apple, when compared to growth in culture, as these genes are most likely to be required for promoting host colonization. For this purpose, we updated the current gene catalogue for isolate MNH120 [19] to increase the total number of annotated genes, including those that encode ECs, which are notoriously difficult to predict in fungi (**Additional file 2: Fig. S1**). In total, 24,502 genes, excluding splice variants, were predicted and, of these, 3,563 were up-regulated and 1,462 were down-regulated at one or more *in planta* time points (*p* value of 0.01 and log2-fold change of 1.5) (**Additional file 3**). It must be pointed out here that our approach was to predict as many genes as possible and, consequently, it is expected that some spurious genes were included in the annotation. However, as many of these spurious genes were anticipated to show a negligible level of expression, most would not have featured in our list of differentially expressed genes, which formed the central focus of our study. For example, of the 24,502 predicted genes, 9,284 had a maximum DESeq2-normalized gene expression count across growth conditions of <1, indicating that they were neither expressed under the conditions tested nor up-regulated *in planta*.

The total set of *in planta* up-regulated genes was used to identify temporal host infection-specific gene expression clusters, henceforth referred to as waves. Here, all expression data were scaled across all samples (Z-score) to visualize the gene expression deviation from the overall mean. For hierarchical clustering, the parameters were set to specifically identify the minimum number of waves for which a distinct gene expression profile could be observed (**Fig. 2A**). In total, five distinct gene expression waves (**Fig. 2A**), representing three separate infection stages (**Fig. 2B**), were identified. More specifically, genes of waves 1 and 2 peaked in expression during early infection at 12 hpi, with expression largely plateauing (wave 1) or trending downwards (wave 2) throughout the remaining infection time points (**Figs. 2A and 2B**). Wave 3 contained genes that peaked in expression during mid infection at 2 dpi (**Figs. 2A and 2B**). Genes of wave 4 displayed their lowest level of expression at 12 and 24 hpi, with expression strongly increasing through 2 and 3 dpi, peaking at 5 dpi during mid-late infection (**Figs. 2A and 2B**). Finally, for genes of wave 5, a similar profile was observed to genes of wave 4, but with expression strongly increasing from 3 dpi and peaking during mid-late infection at 7 dpi (**Figs. 2A and 2B**).

**Fig. 2.**
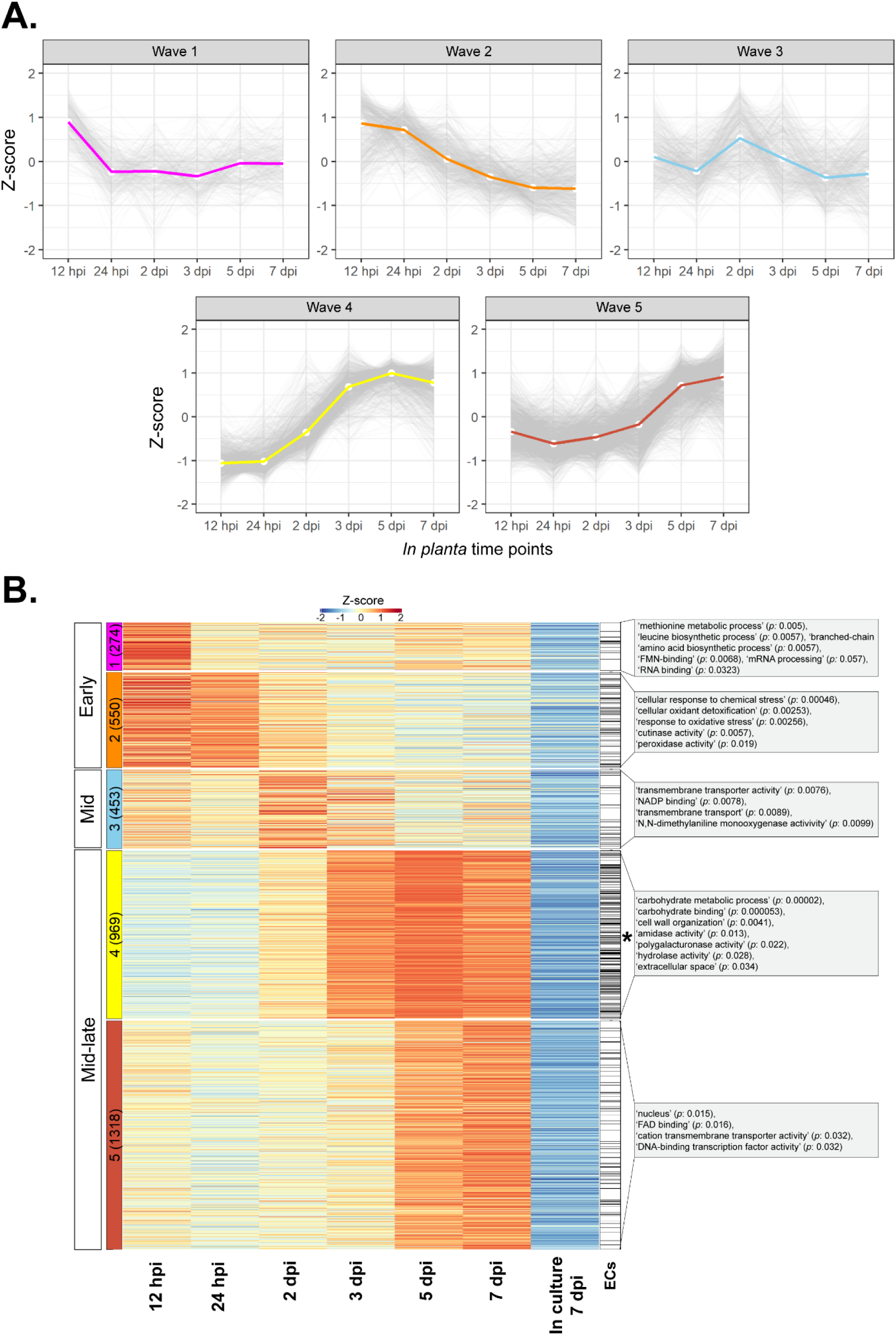
Genes of *Venturia inaequalis* up-regulated during infection of susceptible apple cultivar ‘Royal Gala’, relative to growth of the fungus in culture on the surface of cellophane membranes overlying potato dextrose agar, belong to one of five distinct temporal waves of expression. **A.** Expression profile of the five distinct temporal expression waves at 12 and 24 hours post-inoculation (hpi), as well as 2, 3, 5 and 7 days post-inoculation (dpi), relative to growth in culture (7 dpi). **B.** Heatmap of all *V. inaequalis* genes up-regulated *in planta* when compared with growth in culture. Gene expression data are scaled rlog-normalized counts across all samples (Z-score), averaged from four biological replicates. Genes up-regulated *in planta* were clustered using hclust according to the Ward.D2 and Euclidean distance methods. Early (12 and 24 hpi), mid (2 and 3 dpi) and mid-late infection (5 and 7 dpi) refer to the stage of infection where peak gene expression was observed. Coloured block labels on the left indicate gene expression waves. Numbers in brackets indicate number of genes per wave. The black asterisk indicates the wave significantly enriched for genes encoding effector candidates (ECs) (*p* value: 3.644e-14). Grey boxes indicate enriched gene ontology (GO) terms. *p*: *p* value.

To determine which biological processes are overrepresented in the five temporal gene expression waves, gene ontology (GO) (**Fig. 2B**) and protein family (Pfam) enrichment (**Additional file 4**) analyses were performed. Genes from waves 1 and 2 were mostly characterized by GO terms associated with high metabolic activity, responses to oxidative stress and cutinase activity. Here, cutinases of carbohydrate esterase family 5 (CE5) were abundant (**Additional file 5: Fig. S2**). In contrast, genes of wave 3 were mostly characterized by GO terms associated with transmembrane transport (**Fig. 2B**). Finally, genes of waves 4 and 5 were mostly characterized by GO terms associated with carbohydrate metabolism and transcription (**Fig. 2B**). In wave 4, for example, a GO enrichment for polygalacturonase activity was observed. This, together with the more general enrichment for carbohydrate metabolism, was supported by the high number of plant cell wall-degrading enzyme (PCWDE)-encoding genes in wave 4, most of which were predicted to encode polygalacturonase enzymes of glycoside hydrolase family 28 (GH28) (**Additional file 5: Fig. S2**).

### Genes encoding non-enzymatic proteinaceous effector candidates of *V. inaequalis* predominantly demonstrate peak expression during the mid-late infection stage of apple

From the 24,502 predicted genes of *V. inaequalis*, 1,955 genes were predicted to encode a secreted protein without a transmembrane domain and, of these, 1,369 were predicted to encode a non-enzymatic effector candidate (EC). Here, only small, secreted proteins of ≤400 amino acid residues in length, as well as larger secreted proteins with an effector prediction by EffectorP v3.0, were considered to be ECs (**Additional file 6: Fig. S3A**). Of the 1,369 EC proteins, 518 were identical to proteins predicted by Deng et al. [19] (**Additional file 6: Fig. S3B**). Sequence similarities within our predicted set of ECs were investigated using BLASTp, with the ECs then grouped into families using spectral clustering. Based on this analysis, 759 of the ECs were grouped into 118 families ranging in size from two to 75 members. Of these, 32 families were expanded with five or more members. Due to the observed similarity in sequence between family members, and the fact that expanded families had previously been predicted by Deng et al. [19], we do not expect the gene models encoding these ECs to be spurious. In contrast to the 759 ECs mentioned above, 610 of the ECs were predicted to be singletons that did not belong to any family (**Additional file 7**). Interestingly, ∼22% of the ECs that belonged to families were encoded by genes that either clustered together or were located in close proximity to each other (i.e. within 10 genes) in the genome of isolate MNH120 (**Additional file 7**).

Based on the differential gene expression analysis presented above, 686 of the 1,369 predicted *EC* genes from *V. inaequalis* were up-regulated *in planta*. To determine when these genes peaked in expression, their expression profile across the five temporal waves during host colonization was investigated. In total, ∼20% of the up-regulated genes peaked in expression during early infection, ∼9% during mid infection, and ∼71% during mid-late infection (**Fig. 3**). In all cases, families made up the majority of ECs in each wave (**Fig. 3**). Most of the *EC* genes that peaked in expression during early infection (121) encoded proteins that lacked predicted functional domains. Exceptions included: (1) a family that encoded proteins with an ‘Egh16-like virulence factor’ domain (PF11327) and had sequence similarity to the appressorium-specific Gas1 effector from the rice blast fungus *Magnaporthe oryzae* [38], hereafter named the Gas1-like family, (2) a family that encoded proteins with a ‘stress-up regulated Nod19’ domain (PF07712), hereafter named the Nod19 family, (3) a family that encoded proteins with a ‘hydrophobic surface-binding protein A’ (HsbA) domain (PF12296), hereafter named the HsbA family, and (4) a family that encoded proteins with a ‘common fold in several fungal extracellular membrane proteins’ (CFEM) domain (PF05730) [39], hereafter named the CFEM family.

While the *HsbA* and *CFEM* families possessed genes that mostly peaked in expression during early host colonization (**Fig. 3**), each of these families also had a smaller number of genes that peaked in expression during waves 4 and 5 of mid-late infection (**Additional file 8**: **Fig. S4**). Given that *HsbA* and *cutinase* genes have been shown to be regulated by the same transcription factor in *Aspergillus nidulans* [40, 41], and that most *cutinase* and *HsbA* genes of *V. inaequalis* peaked in expression during wave 2 of the early infection stage (**Fig. 3**, **Additional file 5: Fig. S2**), we set out to determine whether the *cutinase* and *HsbA* genes of *V. inaequalis* were co-expressed. Based on the Pearson correlation coefficient, which was calculated between the *cutinases* and *HsbA* gene expression profiles during the early infection stage, the *cutinase* and *HsbA* genes were indeed found to be co-expressed (R > 0.8, *p* < 0.01). Another family with members exhibiting different expression profiles during host colonization was the *Cin1* family (**Additional file 8**: **Fig. S4**), which is specific to the *Venturia* genus [42]. This family contains the *Cin1* gene (*g8385*), which encodes a cysteine-rich protein with eight repeats, and two *Cin1-like* genes, *Cin1-like 1* (*g10529*) and *Cin1-like 2* (*g13013*), which encode smaller proteins with only one repeat. *Cin1* peaked in expression during wave 4, and was the most highly expressed gene during mid-late host colonization. In contrast, the *Cin1-like* genes peaked in expression during wave 2 of early infection (**Additional file 8**: **Fig. S4**).

**Fig. 3.**
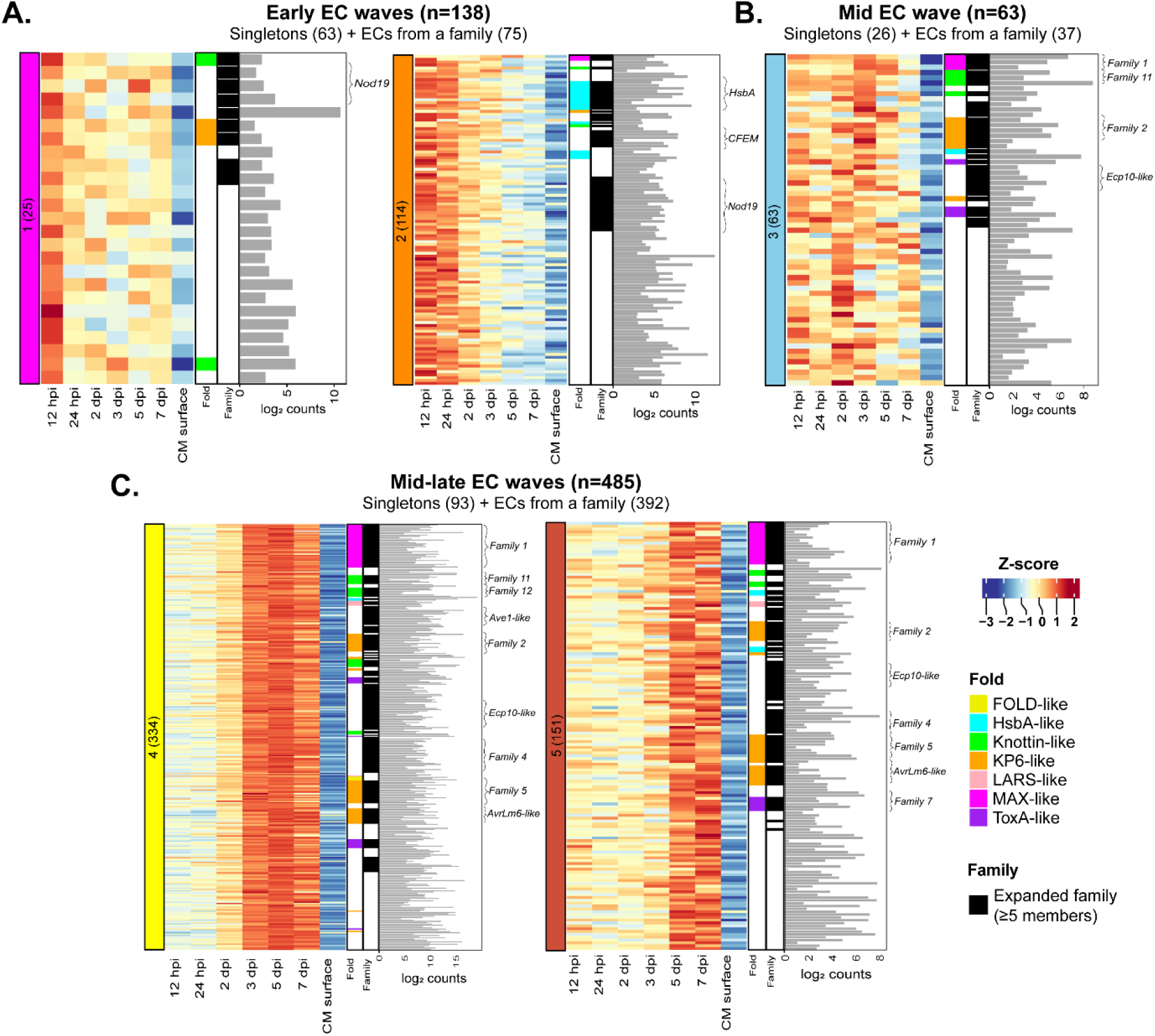
Genes encoding secreted, non-enzymatic proteinaceous effector candidates (ECs) of *Venturia inaequalis* are expressed in temporal waves during colonization of susceptible apple cultivar ‘Royal Gala’. **A.** Heatmap of genes demonstrating peak expression during waves 1 and 2 of the early infection stage at 12 and 24 hours post-inoculation (hpi). **B.** Heatmap of genes demonstrating peak expression during wave 3 of the mid infection stage at 2 and 3 days post-inoculation (dpi). **C.** Heatmap of genes demonstrating peak expression during waves 4 and 5 of the mid-late infection stage at 5 and 7 dpi. Heatmap gene expression data are scaled rlog-normalized counts across all samples (Z-score). Bar plots depict the maximum log2 DESeq2-normalized count value across all *in planta* time points. Black annotations indicate genes encoding proteins that belong to an expanded EC family (≥5 members), with brackets highlighting large families and families with sequence similarity to avirulence (Avr) effector proteins from other plant-pathogenic fungi. CM: cellophane membrane.

The mid and mid-late waves were characterized by 513 genes that mostly encoded EC proteins without a functional domain. The main exception was for members of the Ave1-like family [19], which had an ‘RlpA-like domain superfamily annotation (IPR036908). Notably, most of the expanded *EC* families that encoded proteins with sequence similarity to effectors or Avr effectors from other plant-pathogenic fungi, such as the *AvrLm6-like* [13] and *Ave1-like* [19] families, displayed a peak level of expression during waves 4 and 5 (**Fig. 3**, **Additional file 9: Table S2**). Other examples included the *Ecp10-like* family, which encoded proteins with sequence similarity to the Ecp10-1 Avr candidate from the tomato leaf mold fungus *Fulvia fulva* (formerly *Cladosporium fulvum*), as well as the *Ecp39-like* family, which encoded proteins with sequence similarity to the *F. fulva* EC Ecp39 [43]. An *Ecp6-like* gene (singleton), which encoded a protein with sequence similarity to the Ecp6 effector from *F. fulva* [19, 44], also peaked in expression during wave 4. Interestingly, most expanded *EC* families that peaked in expression during waves 4 and 5 encoded proteins that lacked sequence similarity to other proteins. Examples included the most expanded *EC* family in *V. inaequalis*, family 1 (**Fig. 4**), as well members from family 2 (**Additional file 15: Fig. S7**). In most cases, only one or a few family members were very highly expressed during host colonization (**Fig. 4**).

**Fig. 4.**
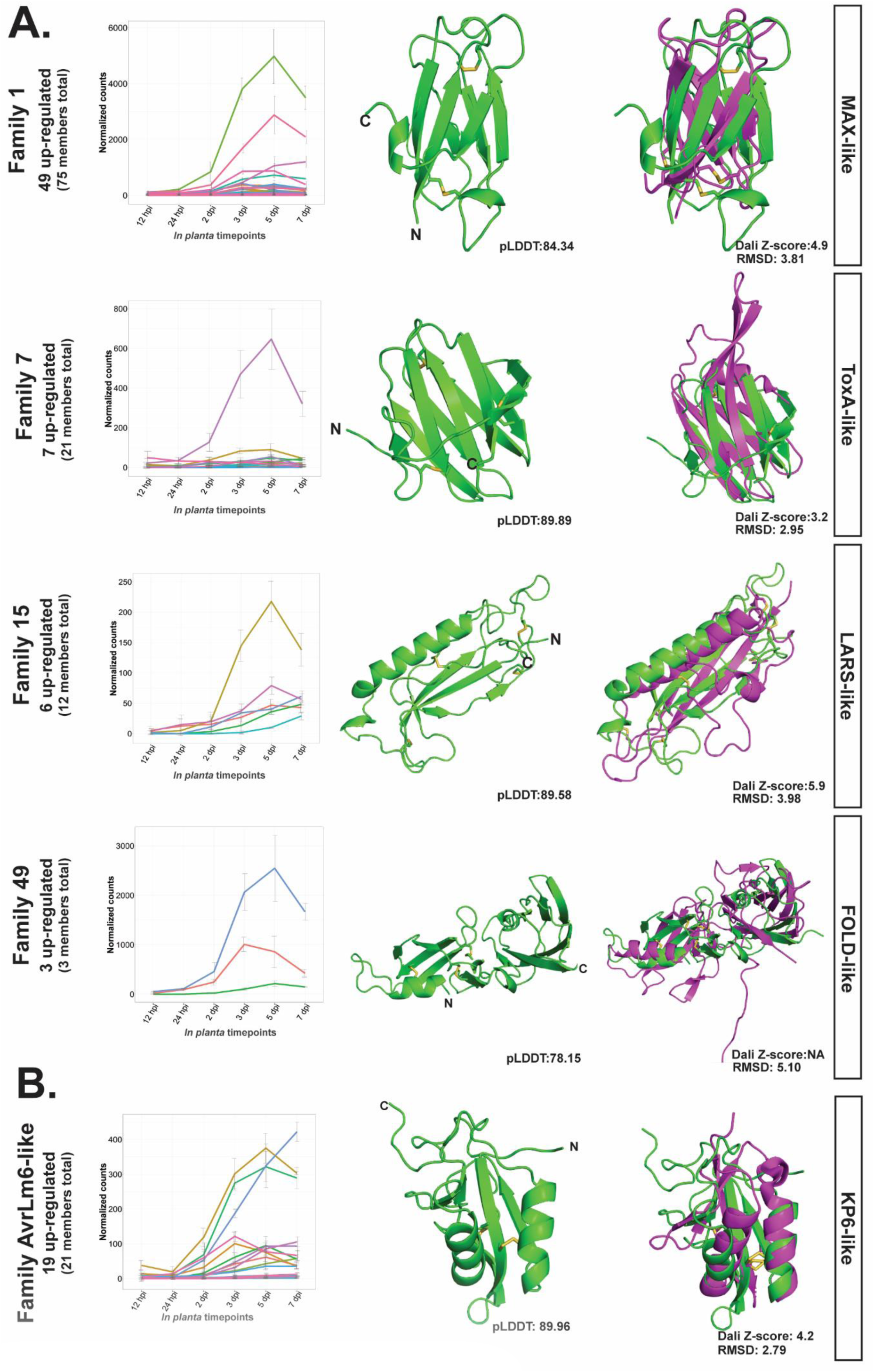
Predicted tertiary structures of secreted non-enzymatic proteinaceous effector candidates (ECs) from *Venturia inaequalis*. **A.** Representative EC family members with structural similarity to avirulence (Avr) effector proteins from other plant-pathogenic fungi. MAX-like: representative predicted family 1 protein structure (green; g13386) aligned to an uncharacterized MAX effector from *Magnaporthe oryzae* (6R5J) (purple). ToxA-like: representative predicted family 7 (green; g4781) protein structure aligned to ToxA from *Pyrenophora tritici-repentis* (1ZLE) (purple). LARS-like: representative predicted family 15 (green; g11097) protein structure aligned to AvrLm4-7 from *Leptosphaeria maculans* (7FPR) (purple). FOLD-like: representative predicted family 49 (green; g3787) protein structure aligned to Avr1/Six4 from *Fusarium oxysporum* (7T6A) (purple). **B**. Predicted EC family members with a KP6-like fold. KP6-like: representative predicted AvrLm6-like family (green; g20030) protein structure aligned to EC Zt-KP6-1 from *Zymoseptoria tritici* (6QPK). Expression data of up-regulated *EC* genes during colonization of susceptible apple cultivar ‘Royal Gala’ are DESeq2-normalized counts, averaged from four biological replicates, with error bars representing standard deviation (hpi: hours post-inoculation; dpi: days post-inoculation). Protein structures predicted by AlphaFold2 represent the most highly expressed member of each EC family from *V. inaequalis*. Disulfide bonds coloured in yellow. N: amino (N) terminus; C: carboxyl (C) terminus. pLDDT: predicted Local Distance Difference Test score (0‒100). A pLDDT score of 70–100 is indicative of medium to high confidence. A Dali Z-score above 2 indicates ‘significant similarities’ between proteins. RMSD: root-mean-square deviation. The protein structure of Avr1/Six4 was not present in the Dali database at the time of writing this manuscript and therefore no Dali Z-score is shown.

Finally, as it known that some fungal effectors are cyclic ribosomally-synthesized and post-translationally modified peptides (RiPPs) called dikaritins [45], and that dikaritin precursor peptides are often mistaken for standard secretory proteins, we set out to determine whether any of our 1,369 ECs were in fact dikaritin precursor peptides. A hallmark of dikaritin precursor peptides is an N-terminal signal peptide followed by multiple perfect or imperfect tandem repeats [46]. As the genes encoding these precursor peptides form part of a biosynthetic gene cluster that also includes a gene encoding a DUF3328 protein, we screened the *V. inaequalis* genome for *DUF3328* genes (and thus, dikaritin gene clusters) to determine which ECs could be dikaritins. In total, nine *dikaritin* gene clusters were identified, with each cluster containing one or more *dikaritin* precursor genes. Based on this analysis, ten of the ECs were identified as putative dikaritin precursor peptides and, of these, four were encoded by genes that peaked in expression during mid-late infection (waves 4 and 5) (**Additional file 10: Fig S5**). The most highly expressed dikaritin precursor gene (*g7830*, from the dikaritin-2 cluster) corresponded to the previously identified gene *Cin3*, which was formerly considered to encode a repeat-containing EC protein [11, 19].

### Several expanded effector candidate families of *V. inaequalis* have predicted structural similarity to Avr effector proteins from other plant-pathogenic fungi

To gain insights into the putative function of ECs, we predicted their tertiary structures using AlphaFold2 [47], and then investigated these structures for similarity to proteins of characterized tertiary structure (and in some cases, function) using the Dali server [48]. This analysis was specifically performed on the most highly expressed member from each EC family (referred to as the representative family member), as well as each singleton, expressed during the temporal host infection-specific waves. In total, the tertiary structure was confidently predicted for the representative family member of 71 (∼76%) EC families and 118 (∼65%) EC singletons (**Additional file 11**).

Strikingly, many EC families were predicted to be structurally similar to one or more ECs or Avr effector proteins with solved tertiary structures from other plant-pathogenic fungi (**Fig. 4**, **Additional file 12: Table S3**, **Additional file 13: Fig. S6**). More specifically, 12 EC families were predicted to be structurally analogous to ECs or Avr effector proteins (**Additional file 12: Table S3**). Remarkably, many of these families were among the most expanded families in *V. inaequalis*. In contrast to the families, only three EC singletons had predicted structural similarity to ECs or Avr effector proteins from other plant-pathogenic fungi (**Additional file 12: Table S3**).

The representative member of the most expanded EC family in *V. inaequalis*, family 1, was confidently predicted to adopt a six-stranded β-sandwich fold with structural similarity to seven ‘*Magnaporthe* Avr and ToxB-like’ (MAX) effectors. The identified MAX effectors were the recently described MAX effector 6R5J [49] and the Avr effectors AvrPiz-t [50], Avr-Pia [51, 52], Avr-Pib [53], Avr1-CO39 [51] and Avr-Pik [54] from *M. oryzae*, as well as the host-selective toxin effector ToxB from *Pyrenophora tritici-repentis* [55], the fungal pathogen responsible for tan spot of wheat (**Additional file 12: Table S3**). A sequence alignment constructed from all family 1 members of *V. inaequalis*, as well as the identified MAX effectors from *M. oryzae*, revealed that these proteins lacked significant sequence similarity, with a maximum pairwise identity of only 13.9% observed between g13386 and Avr1-CO39 (**Additional file 14**). However, these sequence-diverse proteins did share the characteristic conserved disulphide bond between β1 and β5 that has previously been reported for MAX effectors [51] (**Fig. 4A**).

Similarly, two expanded families, families 7 and 28, as well as family 38 and a singleton, were predicted to share a common β-sandwich fold with structural similarity to the host-selective toxin effector ToxA from *P. tritici-repentis* [56], the Avr effector Avr2/Six3 from the wilt fungus *Fusarium oxysporum* [57, 58], and the AvrL567 (AvrL567-A and/or AvrL567-D) Avr effectors from the flax rust fungus *Melampsora lini* [59, 60] (**Fig. 4A**, **Additional file 12: Table S3**, **Additional file 13: Fig. S6**). Notably, despite sharing an overall β-sandwich fold with similar topology, these proteins were very diverse at the amino acid level, with a maximum sequence identity of 12.7% (**Additional file 14**). All of the ToxA-like families had a different number of β-sheets (**Additional file 13: Fig. S6**).

Another two families, families 15 and 47, were predicted to have structural similarity to the AvrLm4-7 and AvrLm5-9 Avr effectors from *L. maculans*, as well as the Ecp11-1 Avr candidate from *F. fulva* [61, 62] (**Fig. 4A**, **Additional file 12: Table S3**, **Additional file 13: Fig. S6**), which belong to the ‘*Leptosphaeria* AviRulence-Supressing (LARS)’ structural effector family [62]. The *V. inaequalis* LARS-like effectors had the same predicted topology as AvrLm4-7 but with a variable number of β-sheets. At the amino acid level, the identified *V. inaequalis* LARS-like proteins shared only 7.8‒16.9% sequence identity with AvrLm4-7, AvrLm5-9 and Ecp11-1 (**Additional file 14**). Interestingly, the *V. inaequalis* structures were predicted to be stabilized by a different number of disulfide bonds and lacked both the conserved disulfide bridge between the α-helix and β-strand, as well as the conserved WR(F/L/V)(R/K) motif, previously reported for the LARS effectors [62].

Finally, family 49 was predicted to adopt a two-domain fold similar to the Avr1/Six4 and Avr3/Six1 Avr effectors from *F. oxysporum* (**Fig. 4A**, **Additional file 12: Table S3**), which are the founding members of the recently identified *Fol* dual-domain (FOLD) structural family of effectors [63]. Despite the *V. inaequalis* FOLD-like family representative sharing only 17.6% amino acid identity with Avr1/Six4 (**Additional file 14**), the cysteine spacing pattern was conserved.

Interestingly, many expanded EC families had a KP6-like fold similar to the KP6 protein of the P6 virus from the corn smut fungus *Ustilago maydis* (**Additional file 15: Fig. S7**) [64, 65] and the Zt-KP6-1 EC from wheat blotch fungus *Zymoseptoria tritici* [66]. This included the AvrLm6-like family, families 2, 5, 23, and 26, as well as two singletons (**Additional file 15: Fig. S7**). All KP6-like proteins were predicted to have between two and four disulphide bridges, two α-helices and a variable number of β-sheets (**Fig. 4B**, **Additional file 15: Fig. S7**).

Unfortunately, many other EC proteins from *V. inaequalis* were too small to be included in RCSB PDB searches (or did not have a significant hit) using the Dali server. To gain further insights into their function, we investigated their general SCOPe fold classification and identified three families and one singleton that were predicted to adopt a knottin-like fold (**Additional file 11**). The most highly expressed member of family 11 (**Additional file 16: Fig. S8**) and a singleton were predicted to adopt a knottin-like fold with two β-sheets and three disulfide bonds that form an intramolecular knot (**Additional file 16: Fig. S8**). Additionally, families 12, 24, and 35 were predicted to adopt a knottin-like fold using the SCOPe database [67] (**Additional file 16: Fig. S8**). However, these proteins were not predicted to have a true intramolecular knot characteristic of knottin proteins based on our current AlphaFold2 predictions. Instead, these proteins shared a common fold with cysteine-stabilized αβ defensins, which are made up of a single α-helix and three β-strands.

In addition to these knottin-like proteins, the Ecp10-like family, along with the candidate Avr effector Ecp10-1 from *F. fulva* [43], were predicted to adopt a small compact β-folded structure that is structurally similar to the PAF protein from *Penicillium chrysogenum* (**Additional file 17: Fig. S9**) [68–70]. Furthermore, the Ecp39-like family, along with the Ecp39 EC from *F. fulva* [43], were predicted to adopt a crambin-like fold (**Additional file 17: Fig. S9**). Unlike that observed for the Ecp10-like family, no analogous protein structures were identified for the Ecp39-like family in the RCSB PDB using the Dali server. Lastly, the Ave1-like family, together with the Ave1 Avr effector from *V. dahliae* [19, 25], were predicted to adopt a double-psi β-barrel fold, stabilized by two conserved disulfide bonds, with high similarity to expansins (**Additional file 17: Fig. S9**).

Lastly, we investigated whether a relationship existed between the predicted fold types of EC proteins and the expression profiles of genes that encoded them. Consistent with the results mentioned above, ECs with a predicted HsbA-like fold were predominantly encoded by genes that peaked in expression during wave 2 of early infection (**Fig. 3**). In contrast, ECs with structural similarity to ECs and Avr effector proteins from other plant-pathogenic fungi predominantly peaked in expression during mid and mid-late infection (**Fig. 3**). Remarkably, all members of the FOLD-like and LARS-like families peaked in expression during mid-late infection (wave 4 and wave 5). Similarly, the majority of members from the MAX-like (65%), ToxA-like (70%), KP6-like (∼59%) and knottin-like (∼66%) families peaked in expression during wave 4.

## Discussion

In this study, we present the first comprehensive transcriptome of *V. inaequalis* during colonization of apple, covering six biotrophic time points from early to late infection. In doing so, we have, to our knowledge, also provided the first comprehensive *in planta* transcriptome of a subcuticular fungal pathogen. Based on this transcriptome, we identified five temporal host infection-specific waves of gene expression for *in planta*-upregulated genes of *V. inaequalis*. Here, genes demonstrated peak expression during one of three stages of host colonization corresponding to early (12 and 24 hpi; waves 1 and 2), mid (2 and 3 dpi; wave 3) and mid-late (5 and 7 dpi; waves 4 and 5) infection. These temporal gene expression waves were biologically distinct from each other and were enriched for different GO terms.

A key focus of our study was to understand the expression profile of *EC* genes that encode secreted, non-enzymatic proteins during host colonization, as these genes make up the bulk of effectors identified from plant-pathogenic fungi to date [4, 6]. Interestingly, during early host colonization, when *V. inaequalis* is either growing on the leaf surface or has just initiated subcuticular growth, only around 20% of the up-regulated *EC* genes peaked in expression. In other plant-pathogenic fungi, however, the percentage of *EC* genes that peak in expression during early host colonization is much higher. For instance, in *U. maydis*, ∼40% of genes encoding secreted proteins were found to be specifically induced during early host-colonization [34]. One possible reason for this difference could be that, during the early infection stage, *V. inaequalis* predominantly colonizes the epicuticular wax above apple epidermal cells, perhaps on its way to colonizing the subcuticular environment. This may suggest that not all infections have yet resulted in close contact with the underlying epidermal cells and, consequently, the mass upregulation of genes encoding effector proteins with roles in suppressing host defences has not yet been initiated.

Among the *EC* genes that peaked in expression during early colonization, several belonged to the *HsbA* family. HsbA proteins have been suggested to recruit cutinases to hydrophobic surfaces [41] and, in *A. nidulans*, *HsbA* and *cutinase* genes are co-regulated by the same transcription factor [40, 41]. Consistent with this previous research, we observed that some *HsbA* and *cutinase* genes of *V. inaequalis* were co-expressed, suggesting that, as previously hypothesized, the HsbA proteins of *V. inaequalis* may recruit cutinases to facilitate the degradation and digestion of the hydrophobic apple cuticle during early host infection [19, 41, 71]. In line with this observation, early infection was enriched for the GO term ‘cutinase activity’ and many CE cutinase-encoding genes were up-regulated. Altogether, these observations support previous reports showing that localized enzymatic hydrolysis is needed for penetration of the apple cuticle by *V. inaequalis* to facilitate access to the subcuticular environment [9, 72].

Other *EC* genes that peaked in expression during early host colonization included members of the *Gas1-like* family, which encode proteins with a ‘Egh16-like virulence factor’ domain, and members of the *Nod19* family, which encode proteins with a ‘stress up-regulated Nod19’ domain. Recently, Gas1-like proteins have been shown to form part of the widely distributed fungal ‘effectors with chitinase activity’ (EWCA) family [73]. EWCA proteins are secreted chitinases without a characterized enzymatic domain in the CAZyme database that degrade immunogenic chitin fragments to prevent chitin-triggered immunity in plants [73]. Thus, it is tempting to speculate that members of the Gas1-like family from *V. inaequalis* play a similar role upon access to the subcuticular environment. The function of proteins with a stress up-regulated Nod19 domain is currently unknown. However, it has been suggested that these proteins are associated with responses to abiotic and biotic stress [74–77]. Based on these studies and the expression of the *Nod19* genes during early infection, the Nod19 family from *V. inaequalis* could play a role in modulating oxidative stress during early subcuticular colonization of apple. Crucially, other genes associated with oxidative stress tolerance, such as those encoding peroxidases, were also enriched during early infection, suggesting that modulation of oxidative stress is crucial for early host colonization by *V. inaequalis*.

Like the *HsbA* family, most members of the *CFEM* family peaked in expression during early host colonization. The CFEM domain is found in several fungal proteins, including those of plant pathogens [78], where it has been shown to confer a diverse range of functions ranging from the promotion or suppression of plant cell death and chlorosis [79–81] to the development of appressoria [82]. Consistent with this functional diversity, and because some members of the CFEM family from *V. inaequalis* instead demonstrated a peak level of expression during mid-late infection, it is likely that the CFEM proteins also play a diverse range of roles in *V. inaequalis* during colonization of apple.

The mid infection stage, which was characterized by the large-scale expansion and continued differentiation of subcuticular infection structures (i.e. stromata and runner hyphae), was enriched for GO terms associated with transmembrane transport. Interestingly, while many *EC* genes of *V. inaequalis* were highly expressed during this infection stage, very few displayed their peak level of expression here. This may suggest that the expression of most *EC* genes is still increasing during mid infection. However, it is important to point out that the mid infection wave was not well defined, overlapping partially with the mid-late infection waves. This is presumably due to the asynchronous nature of *V. inaequalis* infection.

Finally, during the mid-late infection stage, when *V. inaequalis* is heavily colonizing the subcuticular space, most *EC* genes peaked in expression. Based on this expression profile, and the fact that their expression steadily ramped up from the onset of the infection process, we believe that these genes likely play a key role in the establishment and maintenance of biotrophy. Intriguingly, many genes encoding GH enzymes associated with the degradation of the plant cell wall, such as pectin-degrading GH28 proteins, also peaked in expression during this infection stage. As nutrients in the subcuticular environment are likely to be scarce throughout host colonization, it is anticipated that *V. inaequalis* meets a portion of its nutritional requirements through the degradation of the pectin-rich layer located between the cuticle and epidermal cells of apple using these enzymes [83]. Related to this, it is well known that fungal GH28 enzymes can be recognized as MAMPs, while host cell wall fragments released as a consequence of GH28 hydrolytic activity can be recognized as damaged-associated molecular patterns (DAMPs), by PRRs, to activate the plant immune system [7]. With this in mind, a subset of the ECs encoded by genes that peaked in expression during mid-late infection may function to suppress plant defences responses initiated by these GH28 enzymes.

Remarkably, many of the *EC* genes that peaked in expression during mid-late infection encoded proteins that belonged to expanded families. Such a phenomenon, where ECs are known to form part of expanded families, has also been observed in other lineage-specific pathogens, including *Blumeria graminis* [84, 85]. Although the relevance of EC family expansion to *V. inaequalis* is not yet well understood, it is anticipated that this expansion facilitates the diversification of effector function and enables the avoidance of recognition by cognate host R proteins [19]. In any case, repetitive elements are expected to play a major role in the expansion process, enabling both *EC* gene duplication and subsequent transposition to other regions of the *V. inaequalis* genome [86, 87]. In line with this, it has previously been shown that *ECs* of *V. inaequalis*, including members of the *Ave1-like* and *AvrLm6-like EC* families, tend to be closely associated with repetitive elements [13, 19]. Moreover, *EC* genes belonging to the same expanded families have been shown to cluster together in the *V. inaequalis* genome (this study), indicating that tandem duplication events might have occurred. To provide further insights into the process of *EC* family expansion in *V. inaequalis*, a chromosome-level genome assembly of this fungus is now required as, due to its highly fragmented nature (1,012 scaffolds) [19], the current Illumina genome provides an incomplete picture of repeat composition and gene clusters.

To gain insights into the function of EC proteins from *V. inaequalis*, we used the *de novo* folding algorithm AlphaFold2 to predict their tertiary structures, as AlphaFold2 has been successfully benchmarked against effectors of characterized tertiary structure from other plant-pathogenic fungi [63, 88]. One of the main limitations when using AlphaFold2 is that proteins with a low number of homologous sequences in public databases normally result in predictions with low confidence scores. In an attempt to overcome this, we generated custom multiple sequence alignments (MSAs) that included the amino acid sequences of all EC family members identified in this study, many of which were not available in public sequence databases, which greatly improved prediction scores (**Additional file 18: Fig. S10**).

Strikingly, many of the EC families, especially the expanded EC families, demonstrated predicted structural similarity to Avr effector proteins from other plant-pathogenic fungi. The biggest of these was the MAX-like family, which had predicted structural similarity to one of the largest effector/EC families from *M. oryzae*, the MAX family [89, 90]. Intriguingly, a recent computational study of secreted proteins from multiple plant pathogens based on AlphaFold2 concluded that the MAX fold was almost exclusive to *M. oryzae*, with a few members of this family also found in the vanilla black spot fungus *C. orchidophilum* [90]. However, here we show that the MAX-like family has undergone massive expansion and diversification in *V. inaequalis*, highlighting the need for more comprehensive sampling of fungal species using AlphaFold2 to better understand the evolutionary origin and distribution of the MAX structural fold.

In *M. oryzae*, Avrs of the MAX effector family are translocated into host cells, where they are recognized by NLR R proteins [50, 53, 91–94]. Of these, Avr-PikD and Avr1-CO39 directly interact with their corresponding NLR R proteins; an interaction mediated through a heavy metal-associated (HMA) domain that is integrated into the R protein itself. Similarly, the MAX effector Avr-Pik binds and stabilizes an independent HMA protein to modulate host immunity [95]. Altogether, these studies suggest that the MAX fold could be well suited to interactions with HMA domains [96, 97]. It is therefore tempting to speculate that ECs of the MAX-like family from *V. inaequalis* are translocated into host cells, where they interact with HMA domain-containing proteins of apple. Certainly, as Rvi15, an R protein of apple that recognizes the AvrRvi15 Avr effector of *V. inaequalis*, is an NLR [98], it seems likely that a subset of Avrs from this fungus are translocated into host cells.

EC families and singletons with predicted structural similarity to ECs and Avr proteins from the ToxA-like family were also identified in *V. inaequalis*. Interestingly, like that observed in *V. inaequalis*, the recent computational study by Seong and Krasileva [90] showed that ToxA-like ECs are greatly expanded in the cereal stem rust fungus *Puccinia graminis*. The same study also showed that a further four species, *F. oxysporum*, *C. orchidophilum*, *V. dahliae*, and *U. maydis*, have members of this family [90]. However, in the case of these four species, only a few members of the ToxA family could be identified. Thus, *V. inaequalis* appears to have one of the largest repertoires of ToxA-like ECs in fungal species investigated to date.

Like the MAX family, some members of the ToxA-like family (Avr2/Six3 and AvrL567) are translocated into plant cells, where they perform their virulence functions and are recognized by their corresponding R proteins [57, 99]. For example, Avr2/Six3 of *F. oxysporum* functions together with another effector, Six5, to facilitate the cell-to-cell movement of effectors by increasing the size exclusion limit of plasmodesmata [100]. Other members, however, are thought to be apoplastic. For instance, an ortholog of ToxA from the wheat blotch fungus *Parastagonospora nodorum* interacts with an integral membrane protein of wheat from the apoplast to facilitate a cell death reaction that involves the intracellular protein, Tsn1 [101]. As all ToxA-like effectors functionally characterized to date display different virulence functions, no insights into the possible function of ToxA-like proteins from *V. inaequalis* can be made. However, as the Rvi6 R protein of apple, which recognizes the AvrRvi6 Avr effector protein of *V. inaequalis*, is an RLP [102], it is possible that these or other ECs of this fungus identified in our study could be recognised as Avr determinants in the subcuticular environment.

Two other structural EC families in *V. inaequalis* were the LARS-like family [62] and the two-domain FOLD-like family [63]. Of note, only one small EC family of *V. inaequalis* was predicted to adopt the FOLD-like fold. However, it must be pointed out that proteins with this fold are known to be difficult to predict [90] and, as a consequence, some members may have been missed. Interestingly, while the precise functions of the LARS and FOLD effector families are currently unknown, both contain members that suppress the host immune response related to the recognition of another member. More specifically, in terms of the LARS family, AvrLm4-7 suppresses immune responses triggered by AvrLm3 and AvrLm5-9 [103, 104], while for the FOLD family, Avr1/Six4 suppresses immune responses triggered by Avr3/Six1 [105]. Together, this suggests that these protein structures may play a role in molecular mimicry to prevent detection [62, 63, 106]. It will be interesting to determine whether similar relationships are observed for the LARS-like and FOLD-like ECs of *V. inaequalis*.

Aside from the folds described above, an intriguing observation was that many of the EC families and singletons from *V. inaequalis* were predicted to adopt a KP6-like or knottin-like fold. The KP6 fold was first described in the antifungal KP6 protein from the P6 virus from *U. maydis* [64, 65] and, in our study, was predicted to be adopted by the AvrLm6-like family as well as AvrLm6 Avr effector protein from *L. maculans*. Notably, this fold is known to be adopted by the EC Zt-KP6-1 (6QPK) from *Z. tritici* [66] and has also been predicted to be adopted by many effectors such as the Avr effector AvrLm10 from *L. maculans* [90, 107], the BAS4 effector from *M. oryzae* [90, 108], the Ecp28 EC family and Ecp29 EC from *F. fulva* [43], and the CbNip1 necrosis-inducing effector from the sugar beet leaf spot fungus *Cercospora beticola* [109]. Even more interesting, as observed in *V. inaequalis*, the KP6-like fold has been predicted to be the most abundant fold in the *M. oryzae* secretome [89, 90] and is widely shared among phytopathogens [90]. Altogether, this suggests that this KP6-like fold is a widely conserved structural family of effectors in different phytopathogens.

In terms of ECs from *V. inaequalis* that had a predicted knottin-like fold, several families and a singleton were identified. Knottins are small, ultra-stable proteins with at least three disulfide bridges that form an intramolecular knot known to provide stability in hostile conditions such as the plant apoplast [110]. Current evidence suggests that multiple fungal effectors adopt this fold. Indeed, the Avr effector Avr9 from *F. fulva* has previously been suggested to adopt a knottin fold, based on NMR [111] and cysteine bond connectivity [112] data. Furthermore, an EC from the poplar rust fungus *Melampsora larici-populina*, MLP124266, which is a homolog of the AvrPm4 Avr effector from *M. lini*, was recently shown to adopt this fold [113]. Other knottin-like families identified in our study appear to adopt a fold resembling defensins, similar to the VdAMP3 effector of *V. dahliae*, which has antifungal activity and facilitates microsclerotia formation [114].

Curiously, the Ecp39-like EC family from *V. inaequalis*, along with the Ecp39 EC from *F. fulva*, were predicted to adopt a crambin fold, which is commonly found in antimicrobial proteins [115], while the Ecp10-like family from *V. inaequalis*, together with the Ecp10-1 Avr candidate from *F. fulva*, were predicted to adopt a fold with similarity to the antimicrobial PAF protein from *P. chrysogenum* [68–70]. The large number of proteins from *V. inaequalis* that are predicted to have structural (KP6-like, knottin-like, Ecp39-like and Ecp10-like) or sequence (Ave1-like) similarity to antimicrobial proteins suggests that a considerable proportion of the ECs from this fungus may be dedicated to antagonistic fungus–microbe interactions to ward off microbial competitors.

It should be noted that as part of our EC prediction pipeline, we also identified four putative dikaritin RiPP precursor peptides that were encoded by genes displaying a peak level of expression during late infection. Dikaritins are a class of cyclic bioactive peptides that have recently been discovered in fungi [116] and have been hypothesized to play a role as effectors in promoting host colonization [6]. In line with this, Victorin, a host-selective dikaritin toxin from the necrotrophic oat blight fungus *Cochliobolus victoriae*, has been shown to be essential for pathogenicity in oat cultivars resistant to the biotrophic crown rust fungus *Puccinia coronata* [45]. The late-expression profile of the *V. inaequalis* dikaritins, together with the finding that many RiPPs from plants have potent antimicrobial activity [117], may suggest that these peptides, perhaps in addition to some of the ECs described above, promote host colonization through the eradication of microbial competitors in preparation for saprobic growth inside fallen leaf litter.

Taken together, our study on *V. inaequalis*, along with previous studies on effector proteins from *M. oryzae*, *F. oxysporum* and other fungi [51, 62, 63, 89, 90], reinforces the idea that many sequence-diverse fungal effectors share common structural folds. This provides weight to the hypothesis that many fungal effector proteins have in fact originated from ancestral folds and suggests that the genes encoding these effectors have evolved through duplication, followed by sequence diversification, to encode sequence-unrelated but structurally similar proteins. Under this hypothesis, the effector proteins have evolved rapidly to a point where almost all sequence similarity, with the exception of residues involved in the maintenance of the overall structural fold, has been lost [51]. It is of course possible, however, that the appearance of common folds, at least in some instances, could be the result of convergent evolution, whereby certain similar folds have evolved independently in different fungi [51].

Specific protein folds may be common across fungal effector proteins as they provide a stable structural scaffold on which surface or loop features can be altered to enable functional diversification [106]. For those Avr effectors that are directly recognized by their corresponding R proteins, it may also be possible that these alterations extend to the evasion of host recognition. Another possibility is that particular structural folds are well suited to certain functions or to interactions with specific host components [106]. Most likely, though, both above-mentioned scenarios are possible [106]. Future research focussing on the finer details of the distribution of structural effector families among both pathogenic and non-pathogenic fungi, and on the functional characterization of members within these families, will shed more light on this intriguing topic.

## Conclusions

In conclusion, we have performed the first comprehensive gene expression analysis of a subcuticular pathogen, with a specific focus on genes encoding non-enzymatic proteinaceous ECs, during host colonization. In doing so, our study provides valuable new insights into the molecular mechanisms underpinning subcuticular host colonization by this largely understudied class of fungi, including *V. inaequalis*. Notably, in conjunction with structural modelling, we have also provided an enriched list of ECs from which effectors and Avr effectors of *V. inaequalis* can be identified and functionally characterized. Such a resource is desperately needed as, to date, there have been no publications reporting the cloning of *Avr* effector genes from this fungus. Once identified, these *Avr* effector genes will enable the real-time detection of resistance-breaking strains in the orchard. Should Avr effectors of *V. inaequalis* belong to expanded protein families, it may then be possible to engineer their cognate R proteins to recognize features common to the structural fold (direct recognition) or to monitor specific host components targeted by multiple members of the Avr family (indirect recognition). Certainly, with the recent development of CRISPR-Cas9 technology in *V. inaequalis* [118], the functional characterization of ECs, and in particular those that form part of expanded EC families, is now possible. Finally, our study has provided further evidence that many sequence-diverse fungal effectors share common structural folds. Given that the genomes of many other *Venturia* species have now been sequenced [19, 22, 119–125], it will be interesting to determine whether specific effector folds are associated with subcuticular growth or the infection of specific host species.

## Materials and methods

### V. inaequalis isolates

*V. inaequalis* isolate MNH120, also known as ICMP 13258 and Vi1 [19, 126], was used for all bright-field microscopy, transcriptome sequencing, gene predictions and AlphaFold2 protein structure predictions in this study. MNH120 is a race (1) isolate, meaning that it can only overcome resistance mediated by the *Rvi1* gene in apple [19]. This is presumably due to a mutated, absent, or non-expressed copy of the corresponding *AvrRvi1* gene (a functional copy of which is rare among isolates of *V. inaequalis* [127]). MNH120 is, however, anticipated to possess a functional copy of all other known *V. inaequalis Avr* effector genes (*AvrRvi2–20*), corresponding to all other known apple *R* genes (*Rvi2–20*) [19]. Gene predictions from *V. inaequalis* isolate 05/172 [21], which is of unknown race status [128], were used to assist the gene predictions in isolate MNH120.

### Growth in culture for RNA sequencing

*V. inaequalis* isolate MNH120 was grown as a lawn culture from conidia on cellophane membranes (Waugh Rubber Bands, Wellington, New Zealand) [129] overlaying PDA (Difco^TM^, NJ, USA) at 20°C for 7 days under white fluorescent lights (4,300 K) with a 16 h light/8 h dark photoperiod. Four culture plates, representing four independent biological replicates, were then flooded with 1 mL sterile distilled water and the fungal biomass was scraped from the membrane surface using a cell spreader. Following this step, fungal suspensions were transferred to independent microcentrifuge tubes, pelleted by centrifugation at 21,000 x g for 1 min, snap frozen in liquid nitrogen, and then ground to a powder in preparation for RNA extraction. The 7 dpi time point was chosen as *V. inaequalis* had produced enough biomass on the surface of cellophane membranes for adequate RNA extraction.

### Plant infection assays for RNA sequencing and microscopy

Seeds from open-pollinated *M.* x *domestica* cultivar ‘Royal Gala’ (Hawke’s Bay, New Zealand), which is a cultivar susceptible to scab disease caused by *V. inaequalis*, were germinated at 4°C in moist vermiculite with 100 mg/ml Thiram fungicide (Kiwicare Corporation Limited; Christchurch, New Zealand) for approximately two months in the dark. Germinated seedlings were planted in potting mix (Daltons^TM^ premium potting mix; Daltons, Matamata, New Zealand) and grown under a 16 h light/8 h dark cycle with a Philips SON-T AGRO 400 Sodium lamp, at 20°C with ambient humidity. Inoculations with *V. inaequalis* isolate MNH120 were performed on freshly un-furled detached leaves from 4- to 6- week old apple seedlings, as described previously [130], with the exception that 5 µl droplets of conidial suspension (1 x 10^5^ ml^-1^) were used to cover the entire leaf surface. At 12 and 24 hpi, as well as 2, 3, 5 and 7 dpi, four infected leaves, each from an independent seedling, were sampled to give four biological replicates. A microscopic evaluation of infection was then performed on harvested tips from these leaves. Here, leaf tips were cleared and stained according to a previously established protocol [131] and then visualised by bright-field microscopy, with images captured using a Leica DFC 295 digital camera and the Leica Application Suite X (LAS X). Immediately following tip harvesting, leaves were snap frozen in liquid nitrogen and then ground to a powder in preparation for RNA extraction.

### RNA extraction and sequencing

Total RNA was extracted from samples of *V. inaequalis* isolate MNH120 grown in culture, as well as infected leaves, using a Spectrum™ Plant Total RNA Kit (Sigma-Aldrich, St. Louis, MO, USA), with DNA subsequently removed using DNase I (Invitrogen^TM^, Thermo Fisher Scientific, MA, USA). RNA concentration and purity were quantified using a Nanodrop ND-1000 Spectrophotometer (NanoDrop Technologies, Rockland, DE, USA), while RNA integrity was assessed on the Agilent 2100 Bioanalyser (Agilent Technologies, Waldbronn, Germany) using an Agilent RNA 6000 Nano Kit in conjunction with Agilent 2100 Bioanalyzer software. Genomic DNA contamination was excluded by visualisation of RNA on a 0.8% agarose gel and absence of polymerase chain reaction (PCR) amplification products specific to the *actin* gene of apple (GenBank Accession <u>OU745002.1</u> [location 20,713,063–20,714,807]; primers RE45 [5––TGACCGAATGAGCAAGGAAATTACT–3–] and RE64 [5––TACTCAGCTTTGGCAATCCACATC–3–]) [132]. Following these quality control checks, total RNA from each of the samples was sequenced on a HiSeq X platform at Novogene (Beijing, China) via the Massey Genome Service facility (Palmerston North, New Zealand; project number MGS00286). Here, only those RNA samples with an RNA Integrity Number (RIN) value of ≥3.5 were sequenced.

### Gene prediction

The genome sequence of *V. inaequalis* isolate MNH120 [19] was downloaded from the Joint Genome Institute (JGI) MycoCosm portal (https://mycocosm.jgi.doe.gov/Venin1/Venin1.home.html) and a new gene catalogue predicted to accommodate those genes that may have been missed in the initial annotation by Deng et al. [19] (summarized in **Additional file 2: Fig. S1**). Here, we combined three sources of information to generate a more complete gene catalogue. More specifically, for the first source of information, the complete set of predicted genes (coding sequences; CDSs) for *V. inaequalis* isolate 05/172 was downloaded from the National Center for Biotechnology Information (NCBI; https://www.ncbi.nlm.nih.gov/nuccore/QFBF00000000.1/) and mapped to the MNH120 genome using GMAP v2021-02-22 [133] to generate a genome annotation file. Isolate 05/172 was used as the source of information for homology-based gene prediction, as it was deemed to have a more complete set of predicted *EC* genes than the previously predicted gene catalogue for isolate MNH120 [19], based on a crude assessment of how many EC family members were present in the non-redundant (nr) protein database using a tBLASTn search. For the second source of information, we used the RNA-seq data generated from isolate MNH120 in this study to predict open reading frames (ORFs), and thus CDSs, from transcript sequences. More specifically, high quality RNA-seq reads (see the RNA-seq read analysis section below) from one biological replicate of each time point of *V. inaequalis* grown *in planta* and in culture were mapped to the MNH120 genome using HISAT2 v2.2.1 [134, 135], with unmapped reads filtered out using SAMtools v9.2.0 [136]. Then, a genome-guided *de novo* transcriptome assembly was performed with the mapped reads using Trinity v2.12.0 [137]. Likely CDSs were identified using Transdecoder v5.5.0 (https://github.com/TransDecoder/TransDecoder), a program that predicts genes from transcript sequences, in conjunction with a minimum ORF length of 50 amino acids. To support gene predictions, a Pfam domain search was performed to ensure that translated ORFs with one or more characterized functional domains were retained. Additionally, to capture as many EC-encoding ORFs as possible, translated ORFs were scanned against a list of EC proteins identified in the previous gene annotation by Deng et al. [19], supplemented with an in-house database of putative MNH120 effector proteins (de la Rosa and Mesarich, unpublished). Here, a BLASTp e-value threshold of ≤0.05 was employed, with proteins identified using this analysis retained as likely CDSs. Finally, for the third source of information, the original gene catalogue predicted for isolate MNH120 by Deng et al. [19] was downloaded from the JGI MycoCosm portal as above. Together, the three sets of CDSs, derived from the three sources of information described above, were then loaded onto the MHN120 genome as different tracks in Geneious v9.05 [138], and manual curation was performed to create an updated gene catalogue based on a consensus prediction for each gene. More specifically, predictions that were identical across at least two of the three sets of CDSs for a given gene were accepted as correct. Likewise, if for a given gene predictions were different across all CDSs, only the CDS supported by mapped RNA-seq reads (intron–exon boundaries) was selected. Finally, when a CDS was only predicted with one method it was accepted as correct.

### Prediction of protein functions

Protein functional domains were predicted using InterProScan v5.51-85.0 in conjunction with the Pfam, HAMAP, MOBIDB, PIRSF, PROSITE and SUPERFAMILY tools [139], while GO class predictions were carried out using Pannzer2 [140]. N-terminal signal peptides were predicted using SignalP v5.0 [141] and transmembrane (TM) domains were predicted using TMHMM v2.0 [142]. CAZymes were predicted using dbCAN2 in conjunction with the Hotpep, HMMER and DIAMOND tools [143]. Only CAZymes predicted with at least two of the three tools were retained for further analysis. Putative PCWDEs were manually identified based on their CAZy classification (http://www.cazy.org) [144], KEGG description and InterPro annotation. *RiPP* gene clusters were manually identified in the *V. inaequalis* MNH120 genome (**Additional file 19: S1 Text**) through the presence of a gene encoding a protein with a DUF3328 domain in close proximity to a gene encoding a dikaritin precursor peptide with an N-terminal signal peptide, followed by one or more perfect or imperfect tandem sequence repeats of at least 10 amino acid residues in length separated by putative kexin protease cleavage sites.

### Prediction of effector candidates and effector candidate families

Small proteins of ≤400 amino acid residues in length with a predicted N-terminal signal peptide, but without a predicted TM domain or endoplasmic reticulum (ER) retention motif (HDEL/KDEL), were annotated as ECs. This list of ECs was supplemented with proteins of >400 amino acids in length with a predicted N-terminal signal peptide, but no predicted TM domain or ER retention motif, provided that they were predicted to be an effector using EffectorP v3.0 [145]. ECs were grouped into protein families using spectral clustering SCPS v0.9.8 [146]. The identified protein families were then manually curated by eye, taking into account conservation of the N-terminal signal peptide sequence, cysteine spacing, as well as conserved functional domains identified with InterProScan v5.51-85.0. To further refine the list of ECs, proteins with an enzymatic annotation by InterProScan v5.51-85.0 were discarded. In cases where only one or two ECs from a family were predicted to have an enzymatic domain, the EC was retained. To determine whether the ECs had sequence similarity to other proteins, a BLASTp analysis using an e-value threshold of 0.05 was performed against the nr protein database at NCBI.

### RNA-seq read analysis

Methods associated with this section are summarized in **Additional file 2: Fig. S1**. As a starting point for the analysis of RNA-seq reads, all unique genes of *V. inaequalis* isolate MNH120 were identified, and any additional genes that were identical in sequence, representing paralogs (or false gene duplications generated as a consequence of incorrect genome assembly by Deng et al. [19]), were masked to avoid the multimapping of RNA-seq reads. Here, a gff3 file containing only the duplicated genes was generated using AGAT [147], with the sequences subsequently masked using BEDTools v2.30.0 (maskfasta) [148]. Finally, a mappability mask was applied to the MNH120 genome to prevent the multimapping of reads to repetitive genomic regions, including those genes of *EC* families that are known to be highly similar in sequence. To generate the mappability mask, the MNH120 genome was fragmented into all possible 150-bp stretches and mapped to the MNH120 genome using the Burrows-Wheeler Aligner (bwa) [149] with a gap open penalty of 3, gap extension penalty of 3 and max mismatch penalty of 3. Finally, the resulting SAM file from was used to generate the mappability mask (http://lh3lh3.users.sourceforge.net/download/seqbility-20091110.tar.bz2). Raw RNA-seq reads were then filtered, in which adapter sequences, as well as reads with >10% Ns or ≥50% low quality (Qscore: 5) bases, were removed, and the final quality of reads was checked using fastQC v0.11.9 [150]. Next, filtered RNA-seq reads from all samples were mapped to the masked MNH120 genome using HISAT2 v2.2.1 [134, 135] and SAMtools v9.2.0 [136] was used to only keep those reads that mapped to the fungal genome. Uniquely mapped reads were counted using featureCounts from SubRead package v2.0.0 to generate a count matrix [151]. Results from all steps of the RNA-seq analysis were aggregated for quality control assessment using MultiQC v1.11 [152].

### Differential gene expression and clustering

The count matrix (see the RNA-seq read analysis section) was imported to R and a differential gene expression analysis was performed with DESeq2 package v1.32.0 [153]. Pairwise comparisons from all samples were performed and genes with a log2fold change in expression of >1.5 and a *p*adj value of <0.01 during at least one *in planta* infection time point, relative to growth in culture, were considered significantly differentially expressed. Here, multiple testing correction was applied using the Benjamini-Hochberg (BH) method of the DESq2 package, in conjunction with a *p*adj value threshold of <0.01. A PCA plot was then generated using the PCA function of the DESeq2 package. Genes that were up-regulated at one or more *in planta* infection time points were selected for hierarchical clustering. For hierarchical clustering, RNA-seq read counts were first normalized using the rlog method from the DESeq2 package and scaled. Hierarchical clustering of the genes was performed using the hclust function according to the Ward.D2 and Euclidean distance methods, with the minimum number of clusters displaying a distinct expression profile during host infection identified in the resulting clustering dendrogram. Using quantitative guidance from the cutree function, the number of clusters was initially set to 10, then systematically reduced due to observed similarity between clusters, to give five distinct clusters with unique expression trends. Visualization of gene expression clusters (waves), with the expression trends plotted, was performed using ggplot2 v3.3.5 [154], while gene expression heatmaps were generated using Complexheatmap v2.9.1 [155]. Pearson correlation coefficients were calculated to investigate the co-expression of genes inside specific clusters.

### GO and Pfam term enrichment analysis

GO predictions from Pannzer2 [140] with a predictive positive value (PPV) >5 were used for a GO term enrichment analysis across the five distinct gene expression waves. The GO term enrichment analysis was performed with topGO package v2.44.0 [156], using the total set of genes employed for clustering as background and a Fisher’s exact test for all GO terms: Biological Process (BP), Cellular Component (CC) and Molecular Function (MF). GO enrichment analysis results were visualized using ggplots2 v3.3.5 [154]. An enrichment test for Pfam domains was performed using a Fisher’s exact test, with all genes targeted in the clustering analysis used as background.

### Structural modelling of protein tertiary structures

AlphaFold2 [47], in conjunction with the ColabFold notebook [157], was used to predict the protein tertiary structures of EC family members from *V. inaequalis* isolate MNH120. Here, only those family members that were encoded by genes up-regulated *in planta*, relative to growth in culture, were used in this analysis, with only the most highly expressed member targeted for prediction (i.e. as a family representative). In each case, the mature amino acid sequence of the EC (i.e. without its predicted N-terminal signal peptide) was used as input. For the Avr1/Six4-like family, the published pro-domain [63] was also removed. For those ECs that had <30 proteins with sequence similarity in the NCBI database, as identified by a BLASTp analysis in conjunction with an e-value threshold of ≤0.05, a custom MSA was generated and used as input. These custom MSAs, which were built with up to 100 protein sequences, depending on how many protein sequences were available, included all mature EC family members that were unique to the updated MNH120 annotation (i.e. that could not be accessed by the AlphaFold2 search algorithms), as well as similar protein sequences identified through the BLASTp analysis at NCBI. To build the custom MSAs, all sequences were aligned using Clustal Omega [158, 159], with the alignments subsequently converted to the a3m format using ToolSeq [160, 161]. The only exception was for members of the FOLD-like family. Here, in an attempt to improve the structural prediction, the *F. oxysporum* Avr1/Six4 and Avr2/Six3 proteins were manually added to the input sequences to generate a custom MSA, even though these proteins were not identified in the initial BLASTp similarity search. For EC singletons, protein tertiary structures were predicted using AlphaFold2 open source code v2.0.1 and v2.1.0 [47], with preset casp14, max_template_date: 2020-05-14, using mature protein sequences as input. Again, only those ECs that were encoded by genes up-regulated *in planta*, relative to growth in culture, were used in this analysis. All predicted protein tertiary structures with a pLDDT score of ≥70 were considered confident predictions. Protein structures with an pLDDT score of 50–60 that also had an intrinsically disordered region predicted with MobiDB-lite [162] or PrDos [163] were also considered confident predictions.

Predicted EC protein tertiary structures were screened against the Research Collaboratory for Structural Bioinformatics (RCSB) PDB database to identify proteins with similar folds using the Dali server [48]. Here, all hits with a Z-score of ≥2 were considered similar. Protein tertiary structures were visualized and aligned using PyMol v2.5, in conjunction with the alignment plugin tool CEalign [164]. To further investigate similarities between protein tertiary structures, TM-align [165] was used to calculate a root-mean-square deviation (RMSD) value. Finally, the general fold of confidently predicted protein tertiary structures was investigated using RUPEE [166, 167] against the SCOPe v2.08 database [67, 168]. Proteins predicted to have a knottin fold in the SCOPe database were assessed using Knotter 3D to determine whether they had a true knottin structure [110].

## Declarations

### Ethics approval and consent to participate

Not applicable.

## Consent for publication

Not applicable.

## Availability of data and materials

The raw RNA-seq data generated in this study, as well as the count matrix and DESEq2-normalized read counts, have been deposited in the NCBI Gene Expression Omnibus (GEO), and are accessible through GEO Series accession number (GSE198244). The *V. inaequalis* MNH120 gene annotations and associated proteins sequences generated in this study, as well as the output of AlphaFold2 (open source or ColabFold) with the PDB files for the predicted ECs tertiary structures are available at zenodo (10.5281/zenodo.6233645).

## Authors’ contributions

CHM, MR, JKB and KMP conceived the project. JKB, CHM and BM conducted plant infections and prepared samples for sequencing. MR, BH, BM and SdlR performed the bioinformatic analyses. CHM, JKB, MPC and REB provided critical input in experimental design and data analysis. MR, CHM, BH and JKB wrote the manuscript. All authors read, revised, and approved the final manuscript.

## Acknowledgements

We acknowledge use of the New Zealand eScience Infrastructure (NeSI) high-performance computing facilities, as well as their technical support and training services. In particular, we thank Dinindu Senanayake for his consulting support on the high-throughput prediction of protein tertiary structures. New Zealand’s national computing facilities are provided by NeSI and are funded jointly by NeSI’s collaborator institutions and through the Ministry of Science & Innovation’s Research Infrastructure programme. URL: http://www.nesi.org.nz. We thank Drs Erik Rikkerink and Jay Jayaraman for critically reviewing the manuscript, and Dr Simon Williams for providing the *F. oxysporum* Avr1/Six4 and Avr3/Six1 protein structure files ahead of public release.

