## Supplementary Data for "The *Venturia inaequalis* effector repertoire is expressed in waves and is dominated by expanded families with predicted structural similarity to avirulence proteins from other plant-pathogenic fungi"

**Additional file 1: Table S1** RNA-seq transcriptome sequencing read statistics from this study. Samples used for the RNA-seq transcriptome sequencing experiment were derived from an infection time course of *Venturia inaequalis* on detached leaves from susceptible apple cultivar 'Royal Gala' at 12 and 24 hours post-inoculation (hpi), as well as 2, 3, 5 and 7 days post-inoculation (dpi), and during growth of the fungus in culture on the surface of cellophane membranes overlying potato dextrose agar at 7 dpi.

| **Sample name** | **Time point** | **Tissue** | **Total number of reads** | **Number of paired reads mapped to the *V. inaequalis* MNH120 genome** | **Overall alignment rate** |
|---|---|---|---|---|---|
| S12V2 | 12 hpi | Apple leaf (*in planta*) | 72,515,775 | 91,069 | 0.13% |
| S12V3 | | | 71,722,781 | 84,531 | 0.12% |
| S12V4 | | | 65,009,930 | 78,419 | 0.12% |
| S12V5 | | | 61,088,214 | 123,211 | 0.21% |
| S24V1 | 24 hpi | | 67,826,021 | 135,704 | 0.21% |
| S24V3 | | | 67,811,505 | 134,126 | 0.20% |
| S24V4 | | | 75,706,837 | 205,908 | 0.28% |
| S24V5 | | | 78,244,648 | 148,265 | 0.20% |
| S2V1 | 2 dpi | | 47,424,653 | 125,061 | 0.27% |
| S2V2 | | | 68,132,940 | 279,525 | 0.42% |
| S2V3 | | | 73,356,846 | 327,163 | 0.46% |
| S2V5 | | | 63,149,333 | 136,993 | 0.22% |
| S3V1 | 3 dpi | | 46,736,839 | 475,239 | 1.05% |
| S3V2 | | | 48,406,231 | 341,262 | 0.73% |
| S3V3 | | | 63,951,888 | 658,625 | 1.07% |
| S3V5 | | | 46,365,679 | 465,485 | 1.04% |
| S5V1 | 5 dpi | | 40,513,300 | 1,439,733 | 3.68% |
| S5V2 | | | 36,798,369 | 494,654 | 1.39% |
| S5V3 | | | 53,569,444 | 2,965,281 | 5.73% |
| S5V4 | | | 46,221,769 | 1,666,765 | 3.73% |
| S7V2 | 7 dpi | | 47,760,909 | 3,482,001 | 7.55% |
| S7V3 | | | 48,892,959 | 2,331,267 | 4.94% |
| S7V4 | | | 47,854,848 | 4,350,665 | 9.43% |
| S7V5 | | | 43,384,995 | 1,988,632 | 4.76% |
| SS1 | 7 dpi | In culture | 24,522,994 | 22,249,229 | 93.77% |
| SS3b | | | 20,754,255 | 18,915,521 | 93.99% |
| SS4b | | | 22,646,155 | 20,670,316 | 93.89% |
| SS8b | | | 26,675,000 | 24,347,911 | 94.34% |

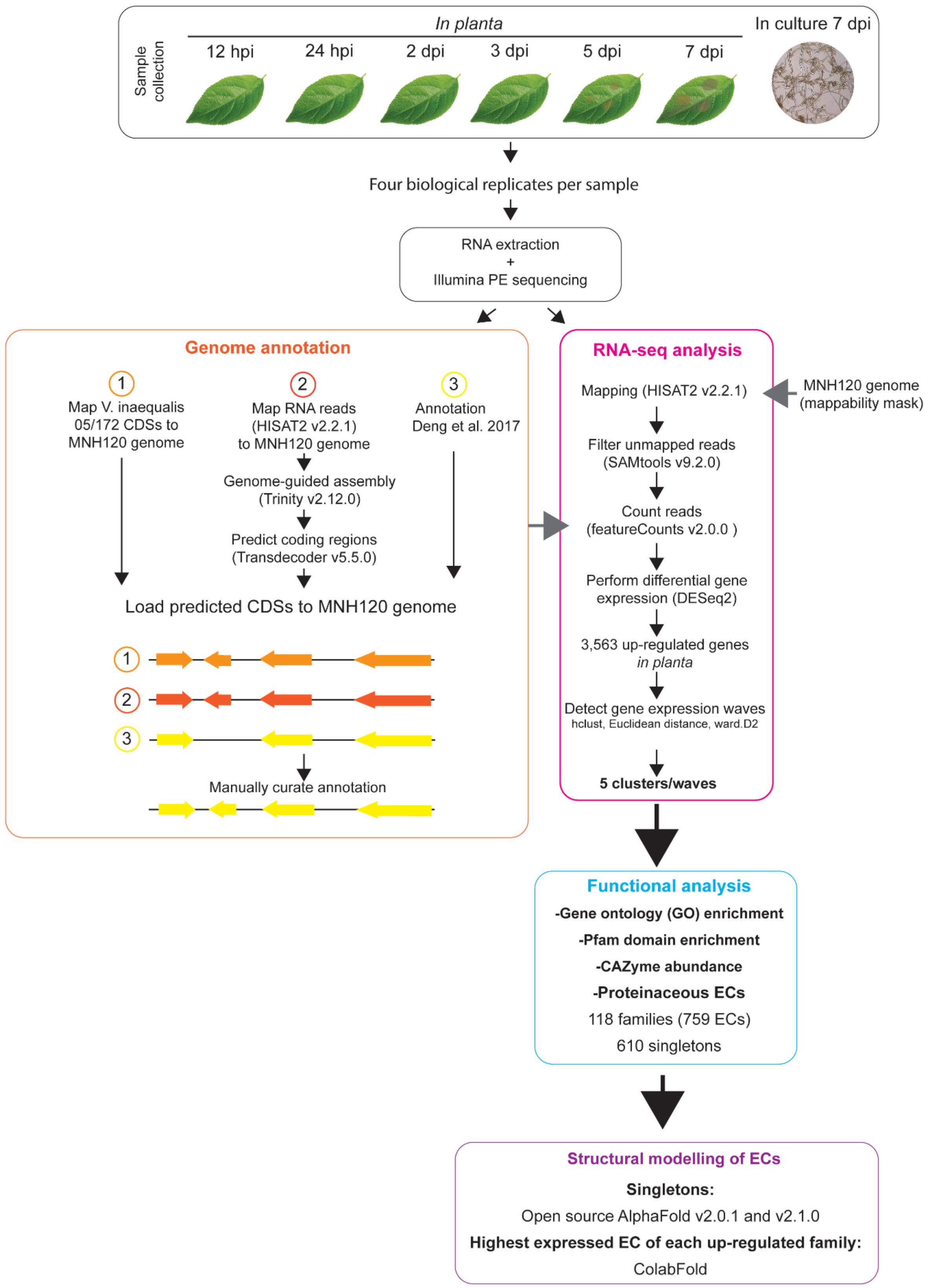


**Additional file 2: Fig. S1** Bioinformatic pipeline used for transcriptome analysis and genome annotation. Total RNA was extracted from apple leaves infected with *Venturia inaequalis* at 12 and 24 hours post-inoculation (hpi), as well as 2, 3, 5 and 7 days post-inoculation (dpi). As a reference for growth in culture, total RNA was also extracted from *V. inaequalis* grown on the surface of cellophane membranes overlaying potato dextrose agar at

7 dpi. Four biological replicates were included per sample. PE: paired-end; CDS: coding sequence; CAZyme: carbohydrate-active enzyme; EC: effector candidate.

**Additional file 3:** Differentially expressed genes of *Venturia inaequalis* during infection of susceptible apple cultivar ‘Royal Gala’, when compared to growth of the fungus in culture on the surface of cellophane membranes overlying potato dextrose agar.

**Additional file 4:** List of Pfam domains in proteins encoded by genes of the five distinct temporal expression waves of *Venturia inaequalis* that are up-regulated during colonization of susceptible apple cultivar ‘Royal Gala’, when compared to growth of the fungus in culture on the surface of cellophane membranes overlying potato dextrose agar.

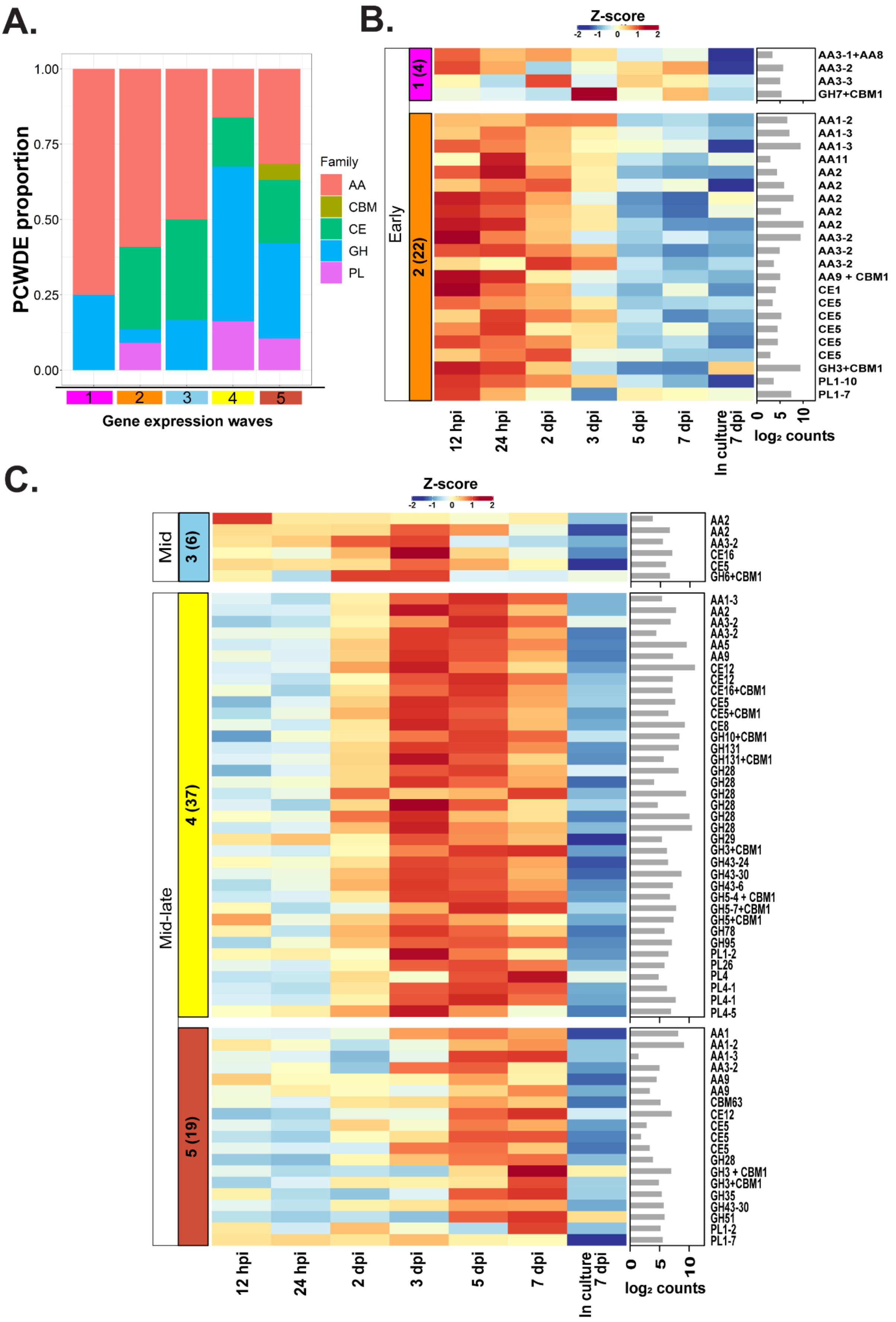

A.
PCWDE proportion
1.00
0.75
0.50
0.25
0.00
Family
AA
CBM
CE
GH
PL
1
2
3
4
5
Gene expression waves
B.
Z-score
-2 -1 0 1 2
Early
1 (4)
2 (22)
AA3-1+AA8
AA3-2
AA3-3
GH7+CBM1
AA1-2
AA1-3
AA1-3
AA11
AA2
AA2
AA2
AA2
AA2
AA3-2
AA3-2
AA3-2
AA9 + CBM1
CE1
CE5
CE5
CE5
CE5
CE5
GH3+CBM1
PL1-10
PL1-7
12 hpi
24 hpi
2 dpi
3 dpi
5 dpi
7 dpi
In culture 7 dpi
0 5 10
log2 counts
C.
Z-score
-2 -1 0 1 2
Mid
3 (6)
AA2
AA2
AA3-2
CE16
CE5
GH6+CBM1
Mid-late
4 (37)
AA1-3
AA2
AA3-2
AA3-2
AA5
AA9
CE12
CE12
CE16+CBM1
CE5
CE5+CBM1
CE8
GH10+CBM1
GH131
GH131+CBM1
GH28
GH28
GH28
GH28
GH28
GH28
GH29
GH3+CBM1
GH43-24
GH43-30
GH43-6
GH5-4 + CBM1
GH5-7+CBM1
GH5+CBM1
GH78
GH95
PL1-2
PL26
PL4
PL4-1
PL4-1
PL4-5
5 (19)
AA1
AA1-2
AA1-3
AA3-2
AA9
AA9
CBM63
CE12
CE5
CE5
CE5
GH28
GH3 + CBM1
GH3+CBM1
GH35
GH43-30
GH51
PL1-2
PL1-7
12 hpi
24 hpi
2 dpi
3 dpi
5 dpi
7 dpi
In culture 7 dpi
0 5 10
log2 counts

**Additional file 5: Fig. S2** Plant cell wall-degrading enzyme (PCWDE)-encoding genes of *Venturia inaequalis* up-regulated during infection of susceptible apple cultivar 'Royal Gala', relative to growth of the fungus in culture on the surface of cellophane membranes overlying potato dextrose agar. **A.** Proportion of *in planta* up-regulated PCWDE-encoding genes in each host infection-specific temporal expression wave. **B.** Heatmap of PCWDE-encoding genes up-regulated *in planta* that demonstrate a peak level of expression during waves 1 and 2 of the early infection stage at 12 and 24 hours post-inoculation (hpi). **C.** Heatmap of PCWDE-encoding genes up-regulated *in planta* that demonstrate a peak level of expression during wave 3 of the mid infection stage at 2 and 3 days post-inoculation (dpi) and waves 4 and 5 of the mid-late infection stage at 5 and 7 dpi. Block labels on the left indicate gene expression wave. Numbers in brackets indicate number of genes per wave. Gene expression data are scaled rlog-normalized counts across all samples (Z-score), averaged from four biological replicates. Labels on the right indicate carbohydrate-active enzyme (CAZyme) classification. Bar plots depict the maximum $log_2$ DESeq2-normalized count value across all *in planta* time points. AA: auxiliary activity; GH: glycoside hydrolase; CE: carbohydrate esterase; PL: polysaccharide lyase; CBM: carbohydrate-binding module.

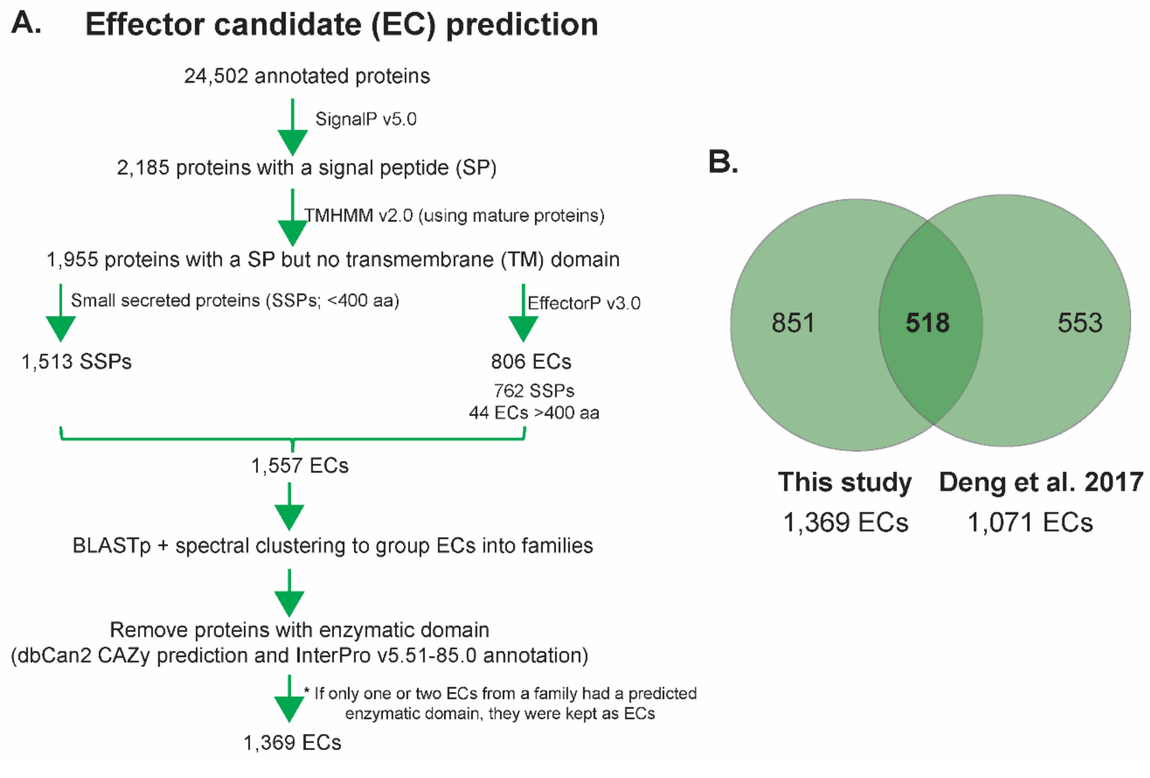


**Additional file 6: Fig. S3 A.** Pipeline for the identification of effector candidates (ECs) from *Venturia inaequalis.* **B.** Comparison of the number of ECs predicted from *V. inaequalis* isolate MNH120 in this study and a previous study (Deng et al., 2017). The comparison is based on an exact protein sequence match. The previous study by Deng et al. (2017) defined ECs as small proteins of <500 amino acid residues in length with a signal peptide and no similarity to lytic enzymes.

**Additional file 7:** List of effector candidates (ECs) identified from *Venturia inaequalis*, their family classification, and indication of whether *EC* family members cluster in the genome.

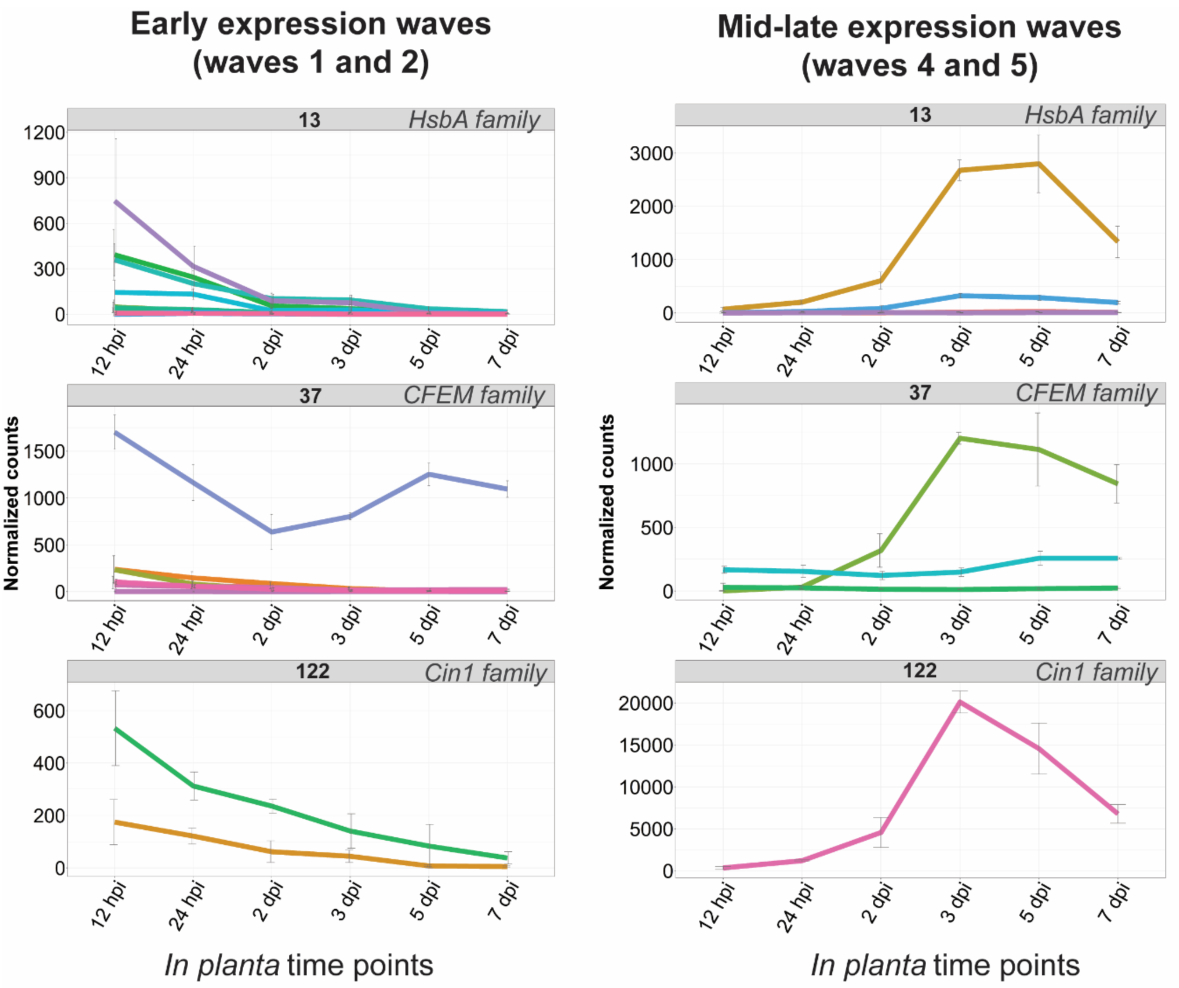


**Additional file 8: Fig. S4** Gene families encoding proteinaceous effector candidates (ECs) of *Venturia inaequalis* that have members demonstrating different expression profiles during early and mid-late colonization of susceptible apple cultivar 'Royal Gala'. Expression data during host colonization are DESeq2-normalized counts, averaged from four biological replicates, with error bars representing standard deviation (hpi: hours post-inoculation; dpi: days post-inoculation). *HsbA*: Hydrophobic surface-binding protein A; *CFEM*: common fold in several fungal extracellular membrane proteins; *Cin1*: Cellophane-induced 1.

**Additional file 9: Table S2** Effector candidate (EC) protein families or singletons from *Venturia inaequalis* that have sequence similarity to EC or avirulence (Avr) effector proteins from other plant-pathogenic fungi.

| EC protein family | Number of family members | Similar EC/Avr and host pathogen | Maximum amino acid identity to similar EC/Avr | Temporal gene expression wave |
|---|---|---|---|---|
| **Gas1-like family** | 2 | *Magnaporthe oryzae* Gas1 | 24.6% | Wave 2 (early infection) |
| **Ave1-like family** | 11 | *Verticillium dahliae* Ave1 | 36.5% | Waves 4 and 5 (mid-late infection) |
| **Ecp39-like family** | 6 | *Fulvia fulva* Ecp39 | 46.9% | Waves 4 and 5 (mid-late infection) |
| **Ecp10-like family** | 61 | *F. fulva* Ecp10 | 23.2% | Waves 4 and 5 (mid-late infection) |
| **AvrLm6-like family** | 31 | *Leptosphaeria maculans* AvrLm6 | 22.7% | Waves 4 and 5 (mid-late infection) |
| **Ecp6-like singleton** | NA | *F. fulva* Ecp6 | 43% | Wave 4 (mid-late infection) |

NA: Not applicable, as the protein is a singleton.

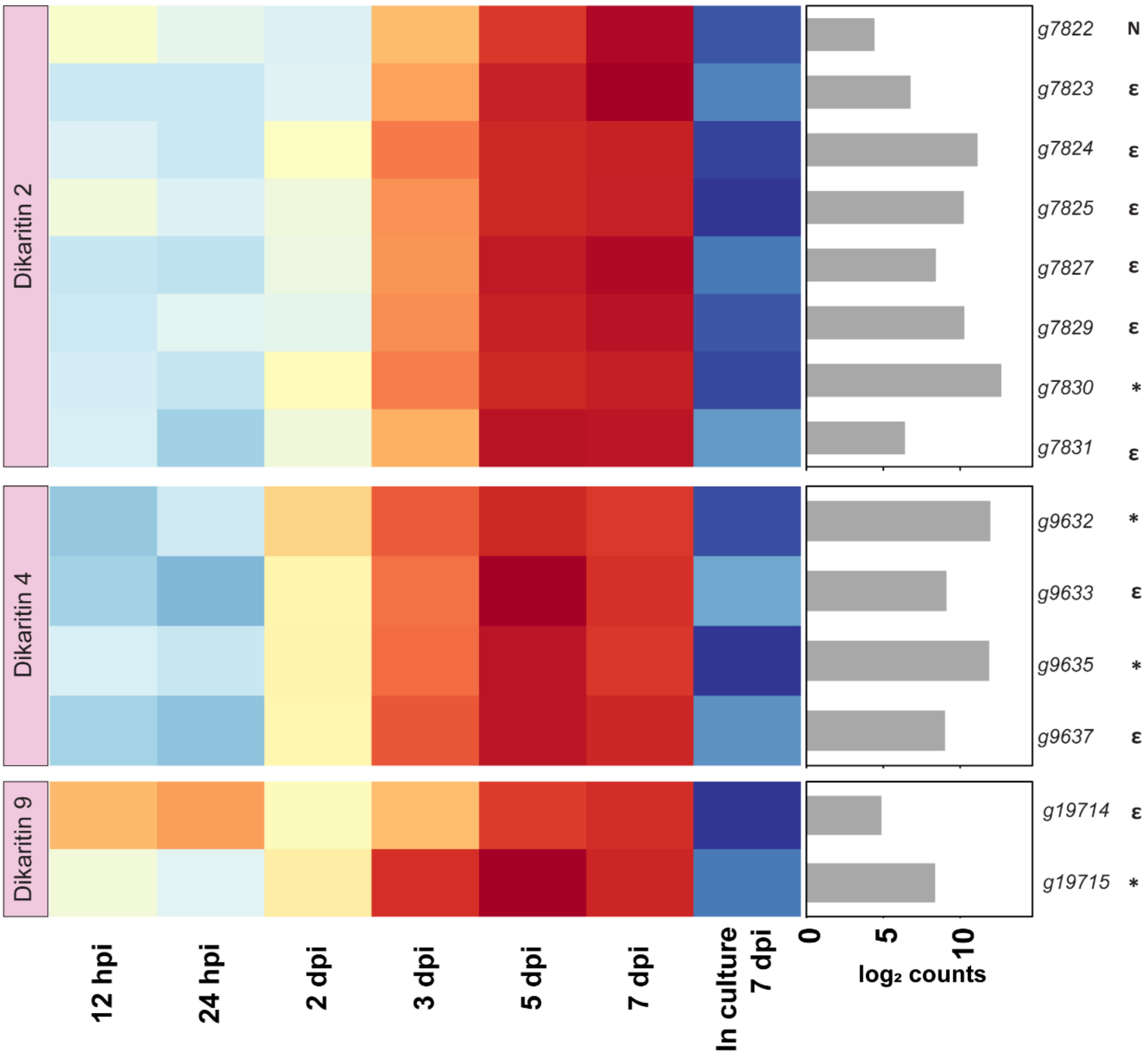


**Additional file 10: Fig. S5** Expression of ribosomally-synthesized and post-translationally modified peptide (*RiPP*) *dikaritin* gene clusters from *Venturia inaequalis* that are up-regulated during colonization of susceptible apple cultivar 'Royal Gala', relative to growth of the fungus in culture on the surface of cellophane membranes overlying potato dextrose agar. Heatmap gene expression data are scaled rlog-normalized counts across all samples (Z-score), averaged from four biological replicates. hpi: hours post-inoculation; dpi: days post-inoculation. Bar plot annotation depicts the maximum $\log_2$ DESeq2-normalized count value across all *in planta* time points. Genes marked as ε putatively encode a protein with a DUF3382 domain or were annotated as a major-facilitator superfamily protein. Genes marked with * putatively encode a dikaritin precursor peptide. The gene marked with N encodes a protein with no characterized functional domain.

**Additional file 11:** List of protein tertiary structures from *Venturia inaequalis* with a confident AlphaFold2 prediction and list of predicted SCOPe folds.

**Additional file 12**: **Table S3** Effector candidates (ECs) of *Venturia inaequalis* (*Vi*) with predicted structural similarity to EC or avirulence (Avr) effector proteins from other plant-pathogenic fungi that have a characterized tertiary structure present in the RCSB PDB.

| *Vi* protein ID | EC family | Number of family members | pLDDT score | Structural similarity (RCSB PDB ID) | Dali Z-score | RMSD |
|---|---|---|---|---|---|---|
| **g13386** | Family 1 | 75 | 84.34 | *Magnaporthe oryzae* MAX effector (6R5J);<br>*M. oryzae* AvrPiz-t (2LW6);<br>*Pyrenophora tritici-repentis* ToxB (2MM2);<br>*M. oryzae* Avr-Pia (2N37);<br>*M. oryzae* Avr-Pib (5Z1V);<br>*M. oryzae* Avr1-CO39 (2MYV);<br>*M. oryzae* Avr-Pik (6FUB) | 4.9;<br>3.7;<br>3.4;<br>3.3;<br>3.3;<br>2.9;<br>2.4 | 3.81;<br>2.72;<br>2.98;<br>3.14;<br>4.71;<br>4.31;<br>3.95 |
| **g11711** | Family 2 | 32 | 97.77 | *Zymoseptoria tritici* Zt-KP6-1 (6QPK) | 5.8 | 2.02 |
| **g18375** | Family 5 | 36 | 89.01 | *Z. tritici* Zt-KP6-1 (6QPK) | 5.7 | 2.34 |
| **g20030** | AvrLm6-like[1] | 31 | 89.96 | *Z. tritici* Zt-KP6-1 (6QPK) | 4.2 | 2.79 |
| **g4577** | Family 23 | 5 | 80.18 | *Z. tritici* Zt-KP6-1 (6QPK) | 5.9 | 3.03 |
| **g12079** | Family 26 | 4 | 88.49 | *Z. tritici* Zt-KP6-1 (6QPK) | 4.9 | 3.15 |
| **g18322** | Singleton | NA | 73.8 | *Z. tritici* Zt-KP6-1 (6QPK) | 4.4 | 3.28 |
| **g4356** | Singleton | NA | 78.02 | *Z. tritici* Zt-KP6-1 (6QPK) | 5.8 | 2.57 |
| **g4781** | Family 7 | 22 | 89.89 | *P. tritici-repentis* ToxA (1ZLE);<br>*Fusarium oxysporum* Avr2/Six3 (5OD4);<br>*Melampsora lini* AvrL567-A (2OPC) | 5.3;<br>3.2;<br>2.7 | 2.95;<br>3.63;<br>3.48 |
| **g13172** | Family 28 | 6 | 59.95* | *P. tritici-repentis* ToxA (1ZLE);<br>*F. oxysporum* Avr2/Six3 (5OD4);<br>*M. lini* AvrL567-D (2QVT);<br>*M. lini* AvrL567-A (2OPC) | 7.4;<br>6.5;<br>5.3;<br>4.9 | 2.47;<br>3.36;<br>3.39;<br>3.78 |

| | | | | | | |
|---|---|---|---|---|---|---|
| **g9034** | Family 38 | 3 | 49.75 | *P. tritici-repentis* ToxA (1ZLE);<br>*F. oxysporum* Avr2/Six3 (5OD4);<br>*M. lini* AvrL567-D (2QVT);<br>*M. lini* AvrL567-A (2OPC) | 5.8;<br>5.0;<br>3.1;<br>2.9 | 3.01;<br>3.28;<br>3.39;<br>3.67 |
| **g4288** | Singleton | NA | 55.02* | *F. oxysporum* Avr2/Six3 (5OD4);<br>*P. tritici-repentis* ToxA (1ZLE);<br>*M. lini* AvrL567-A (2OPC) | 4.5;<br>3.6;<br>2.8 | 3.42;<br>3.38;<br>3.05 |
| **g11097** | Family 15 | 12 | 89.58 | *Leptosphaeria maculans* AvrLm5-9 (7AD5);<br>*L. maculans* AvrLm4-7 (4FPR);<br>*Fulvia fulva* Ecp11-1 (6ZUQ) | 6.5;<br>5.9;<br>5.4 | 3.50;<br>3.98;<br>3.63 |
| **g24490** | Family 47 | 2 | 75.85 | *L. maculans* AvrLm4-7 (4FPR);<br>*L. maculans* AvrLm5-9 (7AD5);<br>*F. fulva* Ecp11-1 (6ZUQ) | 4.1;<br>3.6;<br>3.5 | 3.86;<br>3.61;<br>3.69 |
| **g3787** | Family 49 | 3 | 78.15 | *F. oxysporum* Avr1/Six4 (7T6A);<br>*F. oxysporum* Avr3/Six2 (7T69) | NA;<br>NA | 5.10;<br>4.00 |

pLDDT: AlphaFold2 predicted Local Distance Difference Test score (0–100). A score of 70–100 is indicative of medium to high confidence. RCSB PDB: Research Collaboratory for Structural Bioinformatics Protein Data Bank. Z-score: A Dali Z-score above 2 indicates 'significant similarities' between proteins. RMSD: root-mean-square deviation (a measure of similarity between protein structures). The smaller the RMSD, the more similar the proteins structures are. NA: Not applicable, structures not available on Dali server database.

[1]: Effector candidate family has similarity to the AvrLm6 protein from *L. maculans.*

*: Indicates that the protein is predicted to have an intrinsically disordered region, which reduces the overall pLDDT score.

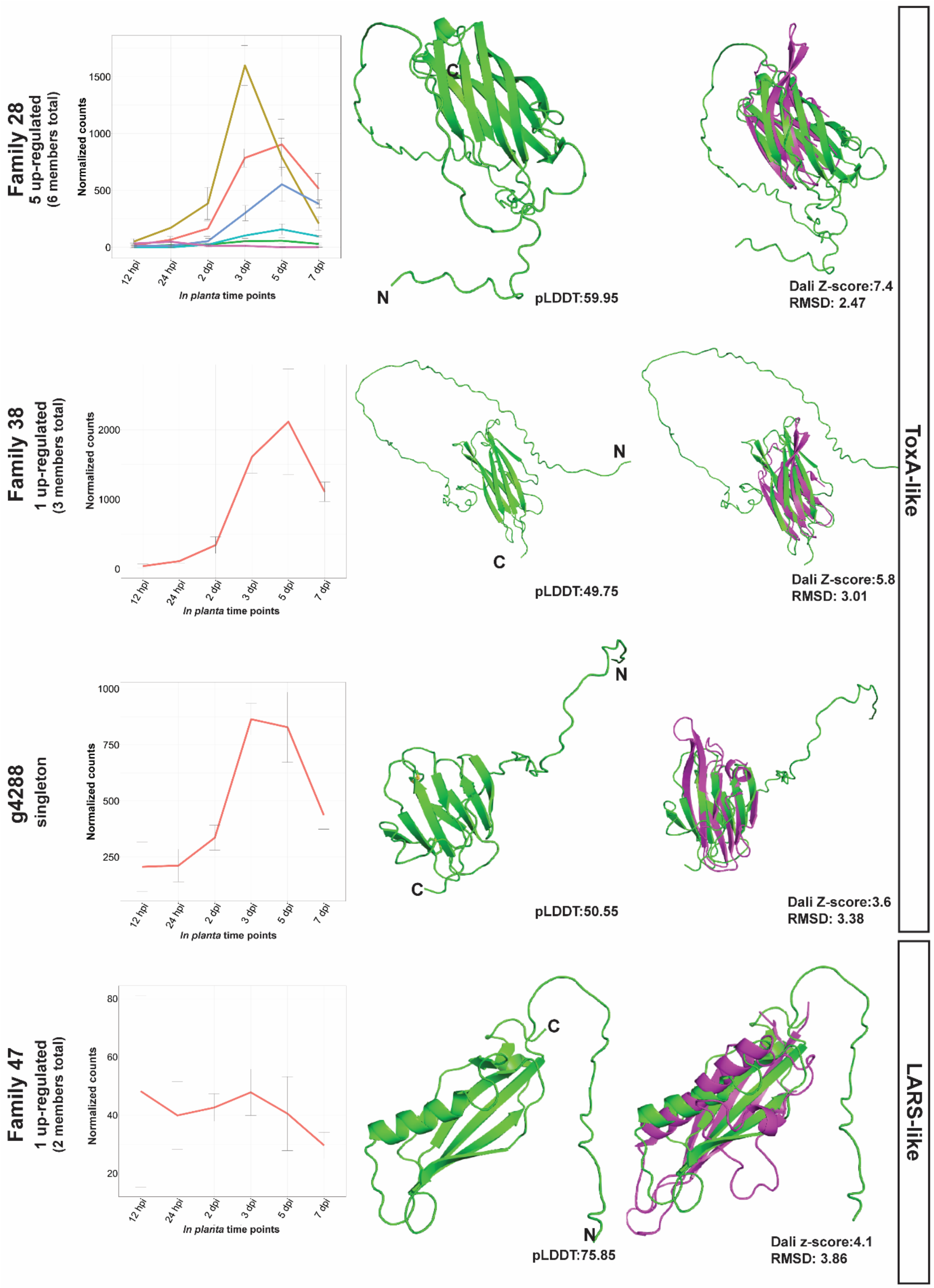


**Additional file 13: Fig. S6** Effector candidates (ECs) from *Venturia inaequalis* with structural similarity to known avirulence (Avr) effector proteins from other plant-pathogenic fungi not shown in Fig 4. Representative *V. inaequalis* protein structures (green; family 28 (g13172), family 38 (g9034), singleton g4288) aligned to ToxA from *Pyrenophora tritici-repentis* (1ZLE) (purple), LARS-like protein structure (green; family 47 (g24490)) aligned to AvrLm4-7 from *Leptosphaeria maculans* (7FPR). Protein tertiary structures predicted by AlphaFold2 are the most highly expressed member from each EC family. Disulfide bonds coloured in yellow. N: amino (N) terminus; C: carboxyl (C) terminus. pLDDT: predicted Local Distance Difference Test score (0–100); a pLDDT score of 70–100 is indicative of medium to high confidence. A Dali Z-score above 2 indicates 'significant similarities' between structures. RMSD: root-mean-square deviation. Gene expression data are from up-regulated *ECs* during host colonization and are based on DESeq2-normalized counts, averaged from four biological replicates, with error bars representing standard deviation (hpi: hours post-inoculation; dpi: days post-inoculation).

**Additional file 14:** Matrix amino acid identity of the effector candidate (EC) structural families (MAX, ToxA, LARS, FOLD).

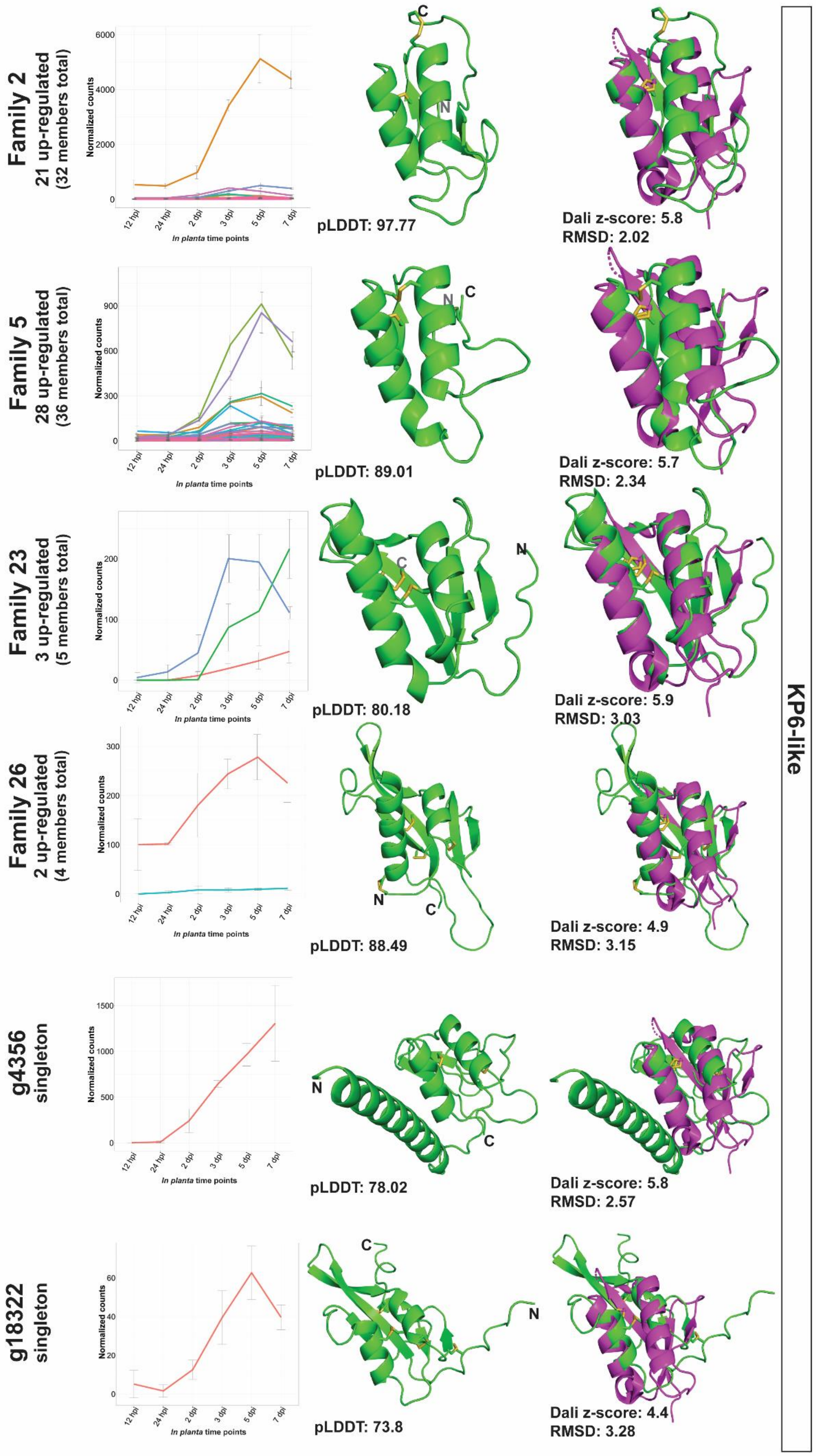

Family 2
21 up-regulated
(32 members total)
pLDDT: 97.77
Dali z-score: 5.8
RMSD: 2.02
Family 5
28 up-regulated
(36 members total)
pLDDT: 89.01
Dali z-score: 5.7
RMSD: 2.34
Family 23
3 up-regulated
(5 members total)
pLDDT: 80.18
Dali z-score: 5.9
RMSD: 3.03
Family 26
2 up-regulated
(4 members total)
pLDDT: 88.49
Dali z-score: 4.9
RMSD: 3.15
g4356
singleton
pLDDT: 78.02
Dali z-score: 5.8
RMSD: 2.57
g18322
singleton
pLDDT: 73.8
Dali z-score: 4.4
RMSD: 3.28
Normalized counts
In planta time points
12 hpi
24 hpi
2 dpi
3 dpi
5 dpi
7 dpi
KP6-like

**Additional file 15: Fig. S7** Effector candidates (ECs) from *Venturia inaequalis* with structural similarity to killer protein 6 (KP6) from *Ustilago maydis* P6 virus not shown in Fig 4. Representative *V. inaequalis* protein structures (green; family 2 (g11711), family 5 (g18375), family 23 (g4577), family 26 (g12079), singleton g4356, singleton g18322) aligned to EC Zt-KP6-1 from *Zymoseptoria tritici* (6QPK). Protein structures predicted by AlphaFold2 are the most highly expressed member from each EC family. Disulfide bonds coloured in yellow. N: amino (N) terminus; C: carboxyl (C) terminus. pLDDT: predicted Local Distance Difference Test score (0–100); a pLDDT score of 70–100 is indicative of medium to high confidence. A Dali Z-score above 2 indicates 'significant similarities' between structures. RMSD: root-mean-square deviation. Gene expression data are from up-regulated *ECs* during host colonization and are based on DESeq2-normalized counts, averaged from four biological replicates, with error bars representing standard deviation (hpi: hours post-inoculation; dpi: days post-inoculation).

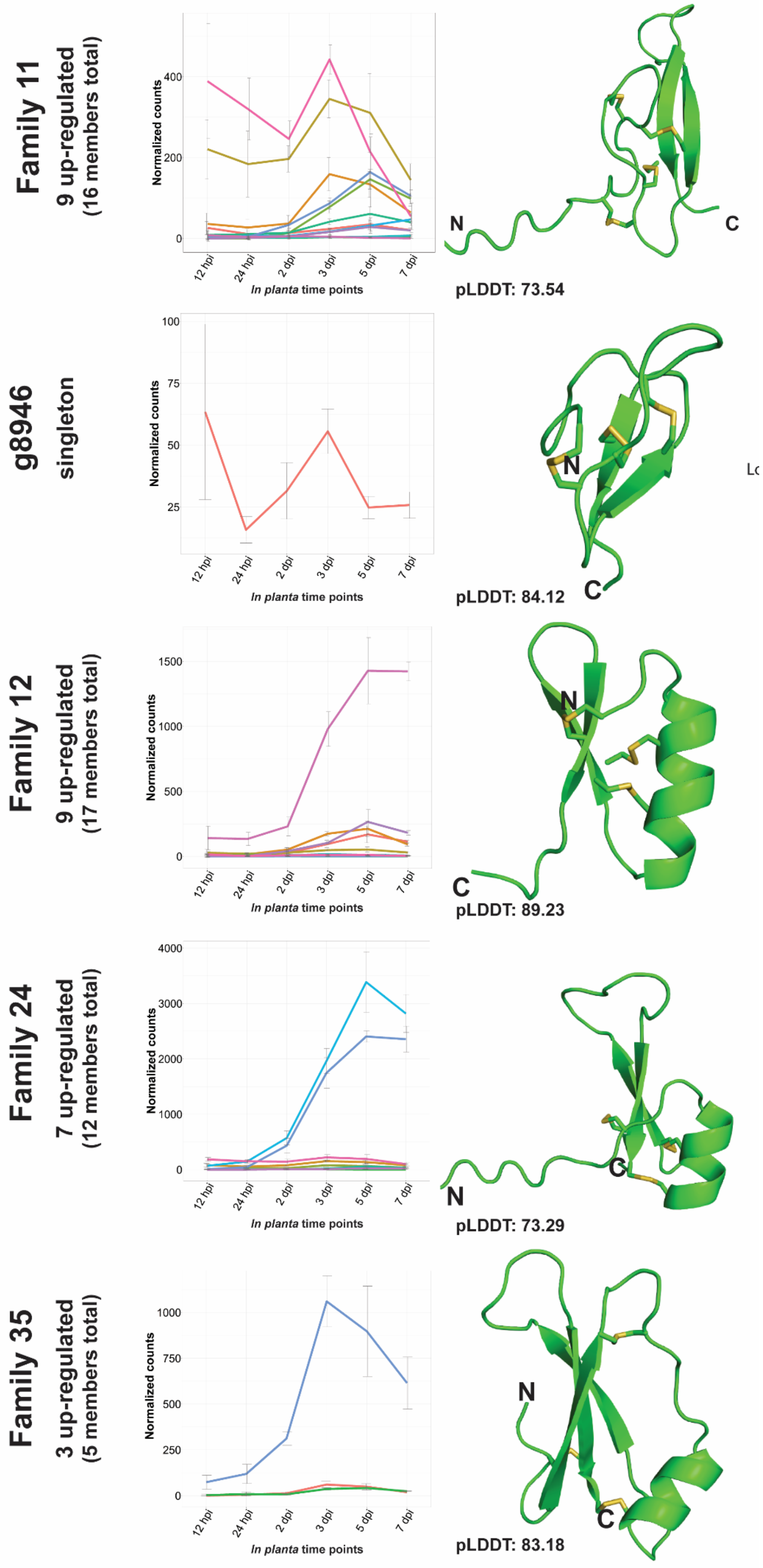
Family 11
9 up-regulated
(16 members total)
pLDDT: 73.54
g8946
singleton
pLDDT: 84.12
Family 12
9 up-regulated
(17 members total)
pLDDT: 89.23
Family 24
7 up-regulated
(12 members total)
pLDDT: 73.29
Family 35
3 up-regulated
(5 members total)
pLDDT: 83.18
Normalized counts
In planta time points
12 hpi
24 hpi
2 dpi
3 dpi
5 dpi
7 dpi
N
C

**Additional file 16: Fig. S8** Effector candidates (ECs) from *Venturia inaequalis* with a predicted knottin-like fold. Representative *V. inaequalis* structures (green; family 11 (g8686), family 12 (g10808), family 24 (g22936), family 35 (g22623), singleton g8946). Protein structures predicted by AlphaFold2 are the most highly expressed member of each EC family. Disulfide bonds coloured in yellow. N: amino (N) terminus; C: carboxyl (C) terminus. pLDDT: predicted Local Distance Difference Test score (0–100); a pLDDT score of 70–100 is indicative of medium to high confidence. A Dali Z-score above 2 indicates 'significant similarities' between structures. Gene expression data are from up-regulated *ECs* during host colonization are based on DESeq2-normalized counts, averaged from four biological replicates, with error bars representing standard deviation (hpi: hours post-inoculation; dpi: days post-inoculation).

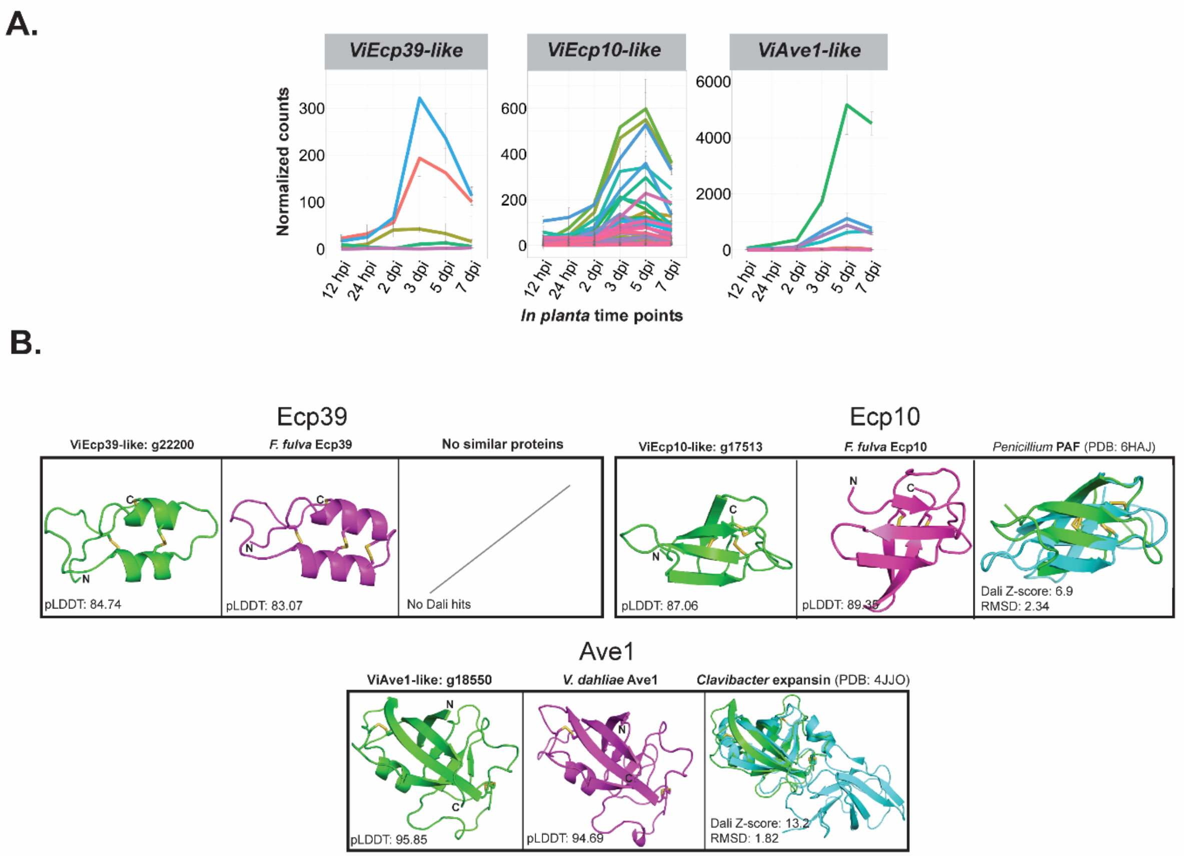


**Additional file 17: Fig. S9** Effector candidate (EC) families from *Venturia inaequalis* (*Vi*) with sequence similarity to effectors and characterized or candidate avirulence (Avr) effector proteins from other plant-pathogenic fungi. **A.** Expression data of *EC* genes during host colonization are DESeq2-normalized counts, averaged from four biological replicates, with error bars representing standard deviation (hpi: hours post-inoculation; dpi: days post-inoculation). **B.** Protein tertiary structures of candidate virulence and avirulence (Avr) effector proteins predicted by AlphaFold2. Disulfide bonds coloured in yellow. *F. fulva* is *Fulvia fulva* and *V. dahliae* is *Verticillium dahliae*. Green tertiary structures represent the *V. inaequalis* protein; purple structures represent the EC/Avr from the other fungal pathogen; cyan, represents closest analogous structure in the RCSB PDB database. Sequence similarity was identified by reciprocal protein searches based on BLASTp (E-value <0.05). pLDDT: predicted Local

Distance Difference Test score (0–100). A pLDDT score of 70–100 is indicative of medium to high confidence. A Dali Z-score above 2 indicates 'significant similarities' between proteins. RMSD: root-mean-square deviation.

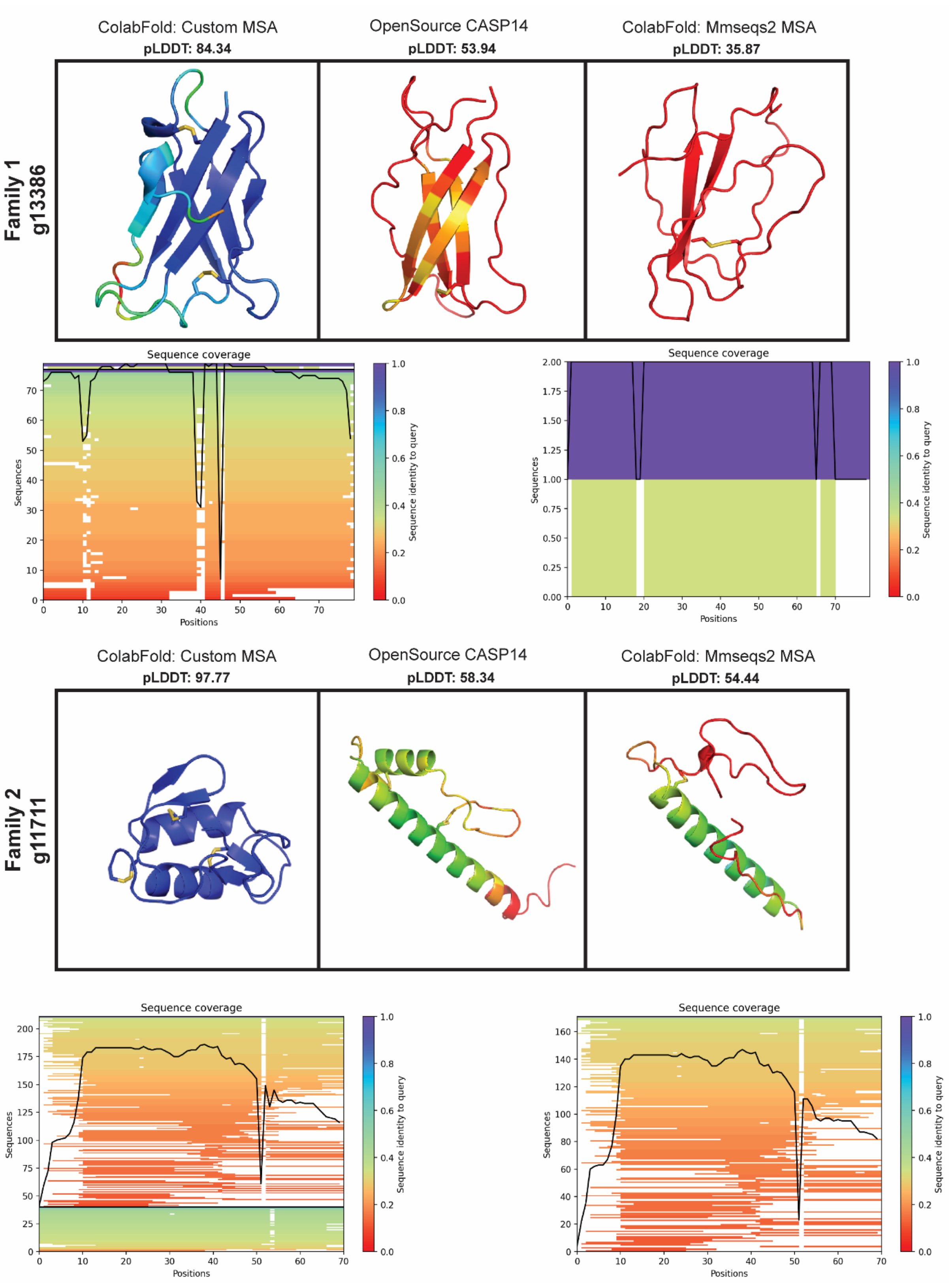

**Additional file 18: Fig. S10** Comparison of protein tertiary structures predicted for two effector candidates (ECs) of *Venturia inaequalis* using different AlphaFold2 methods. EC protein tertiary structures are coloured by amino acid pLDDT (predicted Local Distance Difference Test) score, with a high pLDDT score coloured in blue and a low pLDDT score coloured in red. ColabFold multiple sequence alignments (MSAs) were generated with Mmseqs2 (Steinegger & Söding, 2017) and MSAs generated by AlphaFold2 open source CASP14 were assembled using HHBlits and HMMER (Johnson et al., 2010; Remmert et al., 2011). Sequence coverage graphs of the MSAs used for the structural predictions were automatically generated by ColabFold (Mirdita et al., 2022).

**Additional file 19: S1 Text. Supplementary methods for the identification of *dikaritin RiPP* gene clusters in *Venturia inaequalis*.**

As a starting point for the identification of *dikaritin RiPP* gene clusters in *V. inaequalis*, genes encoding DUF3328 proteins were identified through a tBLASTn analysis of the MNH120 genome in Geneious v9.1.8, using the g7827 DUF3328 protein of *V. inaequalis* (Dikaritin cluster 2) as a query. This was followed by exhaustive reciprocal tBLASTn analyses of the identified sequences against the *V. inaequalis* genome until no new sequences were identified. The resulting protein sequences were then analysed using InterproScan v5.51-85.0 to identify those that carried a putative DUF3328 domain (Jones et al., 2014). Notably, some proteins were not predicted to contain a DUF3328 domain or were instead predicted to belong to the Major-facilitator superfamily. These proteins were manually investigated and proteins with a characteristic DUF3328 domain structure, consisting of a short amino (N)-terminal tail (34–58 aa), a transmembrane (TM) domain (19–24 aa) and a domain containing the tandem HxxCH DUF3328 active site motif, were manually annotated as putative DUF3328 proteins. Up to 10 genes up- and down-stream of the putative DUF3328-encoding gene in the MNH120 genome were then assessed for signatures of a classical dikaritin RiPP precursor-encoding gene (i.e. a gene that encodes a precursor peptide with an N-terminal signal peptide, followed by one or more perfect or imperfect tandem sequence repeats of at least 10 amino acids in length that are separated by putative kexin protease cleavage sites), and analysed for functional domains using InterproScan v5.51-85.0. Finally, to define the borders of *dikaritin RiPP* gene clusters, transcriptome data from this study were used to investigate the expression of genes encoding the DUF3328 domain-containing protein and putative precursor protein, as well as the surrounding genes. Only those genes with a similar *in planta* expression profile to the genes encoding the DUF3328 domain-containing protein and putative precursor peptide were considered to form part of the *dikaritin RiPP* gene cluster in question.
